# How spiders use dragline silk to control abseiling and jumping

**DOI:** 10.64898/2026.09.29.755426

**Authors:** Petur Bryde, Paul Shamble, L. Mahadevan

## Abstract

Spiders arrest their fall by abseiling on dragline silk, varying silk release rate to avoid excessive strains that could break the thread. To understand how this is achieved, we develop a minimal dynamical model of abseiling and formulate a time-optimal control problem subject to physical limits on silk strain and dragline spinning rate. We find that the optimal trajectory includes an interval over which the strain is constant, resulting in uniform deceleration consistent with published experiments. To test the generality of this strategy, we then turn to a distinct extreme motor behavior seen in jumping spiders: using a dragline anchored at the takeoff site to correct their orientation while jumping at speeds exceeding 100 body lengths per second. We show that the same constant-strain strategy suffices to counteract the initial body rotation and ensure an upright landing without significant loss of range. Together, our theoretical framework suggests how strain regulation may serve as a strategy for controlling extreme movements such as abseiling seen in many spiders and may be specialized to tasks such as aerial braking and reorientation in jumping spiders.

## 1. Introduction

A characteristic feature of spiders is their ability to produce silk threads from spinnerets, organs located near the posterior end of the abdomen. Several distinct types of silk have evolved, with material properties suited to different purposes. Dragline silk, formed in the major ampullate gland, is notable for its exceptional tensile strength and extensibility, making it among the toughest naturally occurring fibers [1]. Spiders utilize dragline silk in many ways, including locomotion (e.g. ballooning [2]), communication [3], and as the primary structural component of spider webs [1]. Most spiders continuously draw out dragline silk as they move, periodically attaching it to the substrate. The silk then acts as a safety line in the event of a fall, allowing the spider to climb back to its previous position (figure 1*a*). Jumping spiders (Salticidae), in particular, anchor themselves in this way shortly before leaping (figure 1*b*). Likewise, spiders deliberately drop on a safety dragline to escape a predator [4].

**Figure 1:**
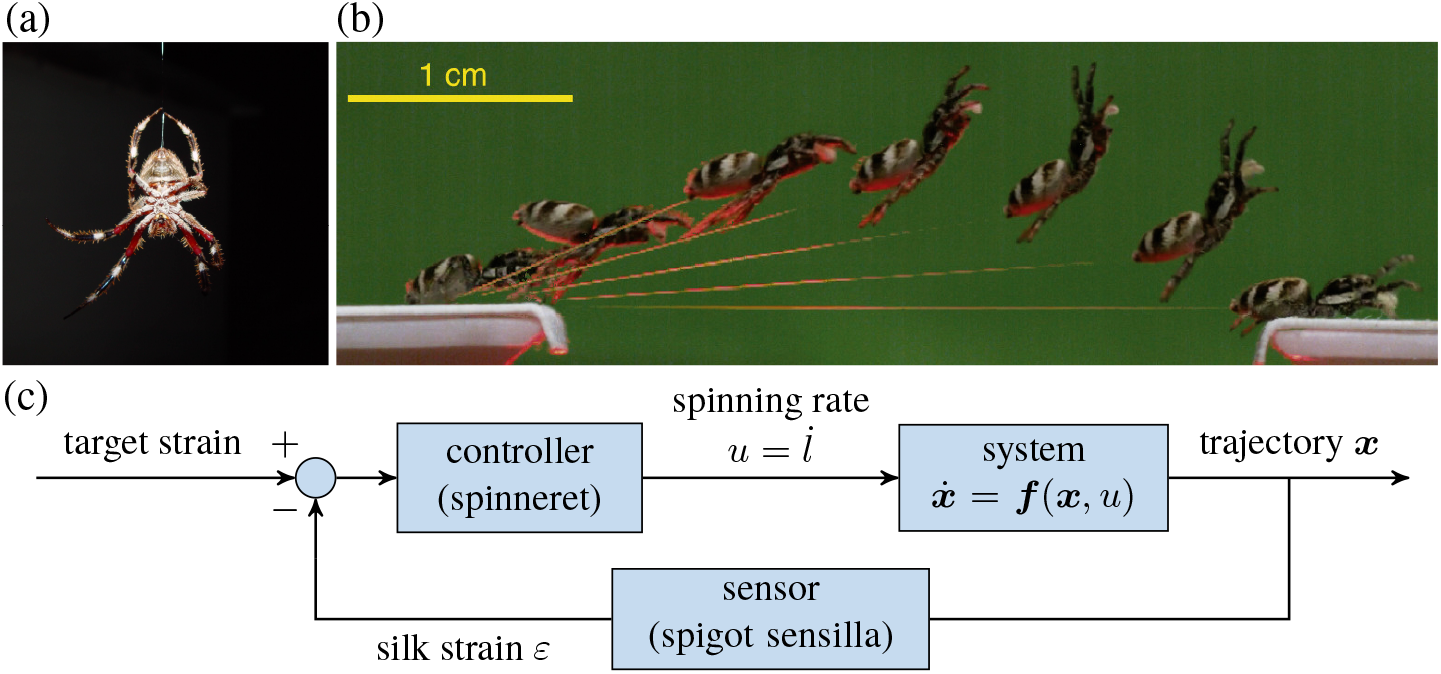
Observations of abseiling and jumping spiders motivate a feedback-control view of dragline silk use. (a) Photograph of a spider suspended from its dragline. (b) Composite image from video (2500 frames/s) of a *Salticus scenicus* jumping across a 3 cm gap, shown at 12 ms intervals; reproduced from Chen et al. [5]. (c) The proposed feedback-control mechanism represented as a block diagram. In the proposed model, silk strain *ε* is sensed and used to regulate the dragline spinning rate 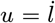 so as to maintain a target strain. The resulting change in silk length affects the trajectory ***x*** through the controlled system 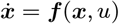. Panel (a): photograph by Leon Brooks, public domain, via Wikimedia Commons.

The process by which spiders arrest their fall using their safety line is more sophisticated than it might first appear. Unlike human bungee jumpers, who are attached to a fixed length of elastic cord, spiders will spool out new silk after the preexisting dragline becomes taut. This behavior may reflect the mechanical limitations of the dragline, which is generally not strong enough to withstand the peak strains that would occur in a bungee jump with a fixed silk length [4,6]. By adjusting the rate at which silk is drawn, the spider can descend on its dragline in a smooth and controlled manner, much like an abseiling climber [4].

Measurements of abseiling spiders indicate that they can precisely regulate their deceleration, and thus the tension in the dragline [7]. Figure 4*d* shows a velocity-time curve for a typical fall by the orb weaver

### Araneus diadematus

Even as the preexisting dragline becomes taut after approximately 5 ms, the spider is essentially in free fall as it continuously spools out new silk. After falling for about 0.13 s, the spider suddenly activates its internal brake and begins to decelerate at a nearly constant rate until coming to rest.

A related braking behavior is observed during the aerial phase of a salticid’s jump. At takeoff, salticids typically acquire a significant rearward rotation, which must be corrected to ensure an upright landing. It appears that they rely on tension in the dragline to control their pitch during flight, and possibly to adjust range and provide stabilization [8,9]. Chen et al. estimated that the jumping spider *Hasarius adansoni* maintains a roughly constant silk force during the initial part of the aerial phase [9]. This suggests a remarkably precise control of silk tension during spooling, even as the spider jumps at speeds near 1 m/s (over 150 body lengths/s) [9]. A separate study of *Salticus scenicus* found that dragline silk produced at the high spinning rates required during jumps, more than ten times typical spinning rates, retains the structural uniformity and high-performance mechanical properties characteristic of silk produced at low speeds [5].

In both cases, it appears that the silk spooling rate is controlled by muscles in the spinnerets, though the exact mechanism of this internal brake is not yet known [10,11]. The precise control exhibited during abseiling and jumping suggests the presence of feedback based on measurements of silk strain. Indeed, certain mechanoreceptors on the spinnerets have been identified that appear to provide information about forces acting on the dragline, and may play a role in regulating silk release [12]. Together, these observations motivate the feedback-control picture shown in figure 1*c*, in which dragline strain is used to regulate the spinning rate and thereby the spider’s trajectory. How strain information is translated into a spooling rate that allows the spider to control its speed or orientation without exceeding the strength of the dragline remains an open question. In particular, it is unclear whether salticids employ a distinct spinning reflex during jumping, or whether the same strain-regulated mechanism governs both behaviors.

Here we attempt to gain insight into these questions using methods of control theory. In section 2, we analyze a simplified model of abseiling and seek control strategies that arrest the spider’s fall without breaking the silk. We first ask whether a simple proportional feedback law based only on silk strain, a quantity the spider can likely sense, can arrest the fall. We show that this feedback law brings the spider to rest asymptotically, but tighter limits on maximum strain lead to slower convergence and greater silk payout. This trade-off motivates a time-optimal control problem subject to constraints on silk strain and dragline spinning rate. We show that, in the biologically relevant regime, time-optimal control predicts a phase of constant silk strain and hence constant deceleration, consistent with behavior observed by Ortlepp and Gosline [7]. Both strategies can be implemented using only measurements of silk strain. In section 3, we show that the same constant-strain strategy can regulate body orientation during aerial motion.

## 2. The abseiling task

### (a) Mathematical model

We consider an idealized model of the abseiling spider as a point mass *m* suspended from a platform by an elastic string representing the silk dragline. We first formulate the model in dimensional variables; whenever dimensional and nondimensional versions of the same quantity are used, we denote the dimensional quantity by a hat 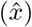 and its nondimensional counterpart by the corresponding unhatted symbol (*x*). Let 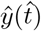 be the vertical distance from the platform at time 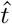, and 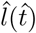 the unstretched length of the silk, as shown in figure 2*a*. We assume that the silk is massless and the strain 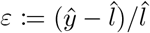 is constant along the dragline. The silk does not resist compression, but exerts a force *F*_*s*_ = *Kmε*^+^ = *Km* max(*ε*, 0) when stretched. Here *K* := *EA/m*, with *E* the silk’s Young’s modulus and *A* its cross-sectional area. This force is counteracted by a gravitational force *F*_*g*_ = *mg*. We neglect air drag, which is small compared to the other forces considered. We take the dragline spinning rate 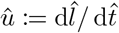 to be the control variable. We assume that *û* is nonnegative and that drawing new silk (û *>* 0) is possible only while the dragline is taut 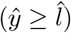.

**Figure 2:**
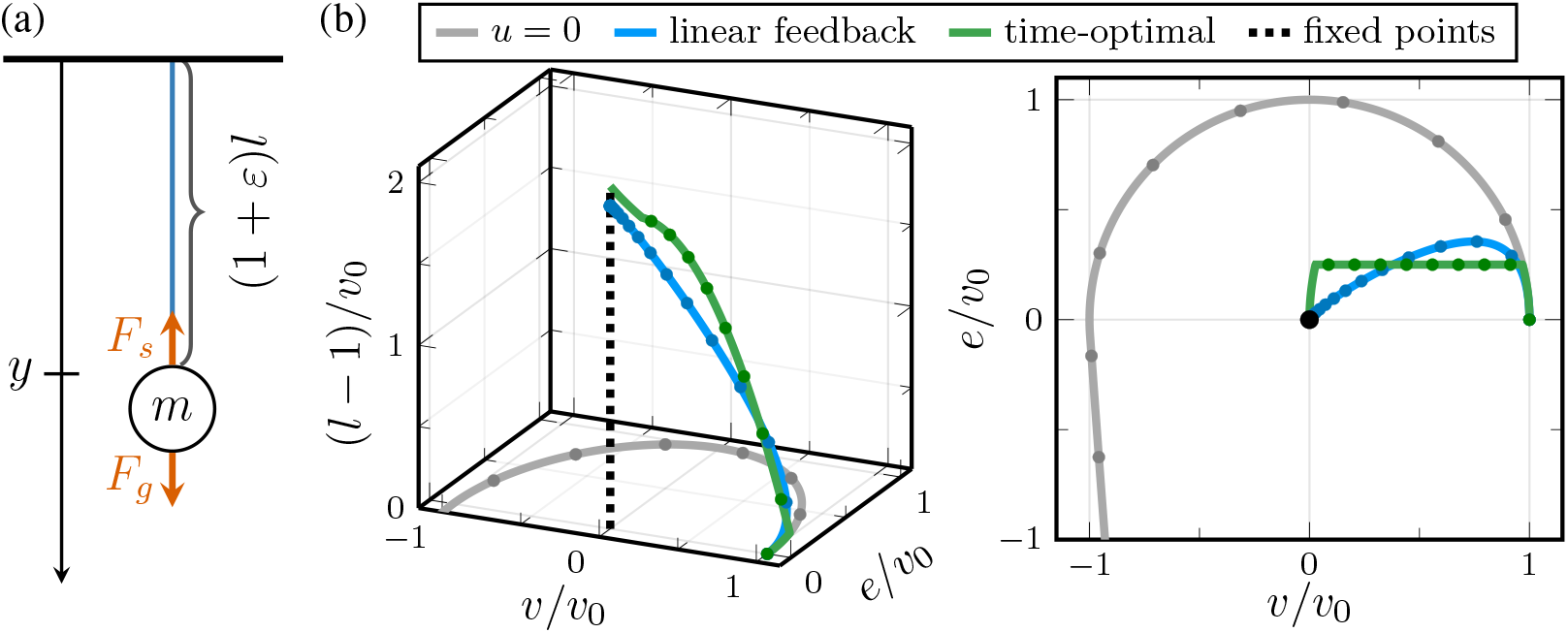
Idealized abseiling model and controlled trajectories. (a) The spider is modeled as a point mass *m* suspended from a platform by an elastic silk thread of rest length *l* and strain *ε*. The dragline spinning rate 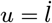 is taken as the control variable. (b) Trajectories of the nondimensional abseiling system, equation (2.2), in the normalized coordinates *v/v*_0_, *e/v*_0_, and (*l*− 1)*/v*_0_, where *v*_0_ = *v*(0), together with their projection onto the (*v/v*_0_, *e/v*_0_)-plane. The gray curve shows the uncontrolled trajectory *u* = 0, corresponding to a bungee jump with fixed silk length. The blue curve shows the trajectory under the feedback law (2.5), with *γ* = 1.0. The green curve shows the time-optimal trajectory subject to *e*≤ *e*_*m*_ = 0.25*v*_0_ and *u* ≤ *u*_*m*_ = 0.2. Markers along the curves are equally spaced in time, showing that the time-optimal trajectory reaches the ray of fixed points in finite time, whereas the feedback trajectory approaches it asymptotically. Both control strategies steer the system to the ray of fixed points *v* = *e* = 0, shown in black, while restricting the excess strain *e*. In contrast, the uncontrolled trajectory misses the ray of fixed points and reaches a larger maximum strain. In all cases, *a* = *g/K* = 0.01. The initial velocity is 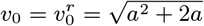, corresponding to a fall starting from rest.

Force balance for the spider yields the equations of motion:

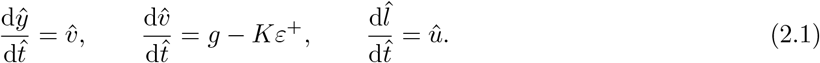

Here 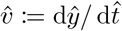 is the velocity of the spider, which is in free fall (*ε <* 0, *F*_*s*_ = 0) until the dragline becomes taut. Eventually, *ε* will increase to the point that *F*_*s*_ = *F*_*g*_, after which the spider will begin to decelerate. We take this moment to be 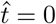 and define 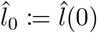 and 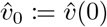.

The *abseiling task* is to choose the spinning rate 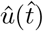 so that the spider comes to a stop, either in finite time or asymptotically as 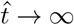, possibly while satisfying constraints on û and *ε*. We say that the spider has stopped if 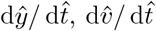, and 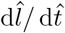 vanish simultaneously; equivalently, if 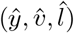 is a fixed point of system (2.1) with û = 0. The fixed points fall on a ray (half-line) characterized by 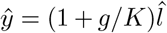,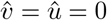, where 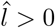 can be arbitrary. If the spider stops at time 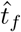, it remains stationary provided 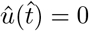for 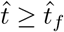.

We nondimensionalize the equations of motion (2.1) by rescaling distances by 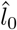 and time by 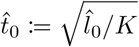, writing 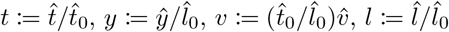, and 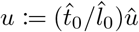. Since the strain is invariant under this rescaling, we have *ε* = (*y − l*)*/l*. Under this rescaling, gravity enters through the dimensionless parameter *a* := *g/K*. Since we are interested in convergence to the ray of fixed points, we make a further change of variables to ***x*** := (*v, e, l*), where *e* := *ε* − *a* is the excess strain. We find that ***x*** satisfies

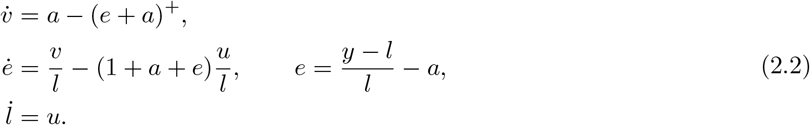

Here and in what follows, dots denote derivatives with respect to the nondimensional time *t*. The initial condition becomes ***x***(0) = (*v*(0), *e*(0), *l*(0)) = (*v*_0_, 0, 1), where 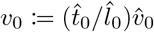. After the change of variables, the ray of fixed points coincides with the positive *l*-axis. We denote this equilibrium set by *E* := {(0, 0, *l*) | *l >* 0}. Measurements of the Young’s modulus and cross-sectional area of dragline silk, together with the spider’s body mass, give typical values of *a* between 0.01 and 0.04 for *Salticus* [5,6], and 0.004–0.01 for *Araneus* [6]. For both species, *a* generally increases with the individual’s body mass. The initial silk length 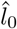 varies depending on the details of the fall, such as how long the spider waits before engaging its friction brake; lengths of order 0.01 m to 0.1 m are typical. For a fall starting from rest at *y* = 0 with *u*(*t*) = 0 for *t <* 0, the initial velocity is 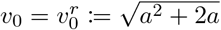, though we allow *v*_0_ to vary in the analysis below. The maximum silk spinning rate is not well characterized and presumably varies between species and, possibly to a lesser extent, between individuals of the same species. We therefore denote the nondimensional maximum spinning rate by *u*_*m*_ and treat it as a model parameter. Observations of abseiling *A. diadematus* have shown that spinning rates exceeding 1 m/s are possible, suggesting an order of magnitude *u*_*m*_ = *O*(10^*−*1^) [7].

Motivated by observations that some spiders decelerate throughout the braking phase until coming to rest [7], we focus primarily on solutions for which *e* remains nonnegative. This is also consistent with our intuition that oscillatory solutions are undesirable. We note that for trajectories satisfying *e* ≥ −*a*, the first equation of system (2.2) becomes 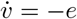. This modified system can be written in the control-affine form:

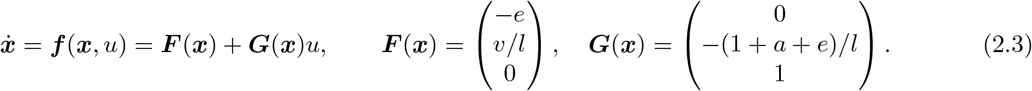

In what follows, we use the simplified system (2.3) for trajectories that remain in the region *e*≥ − *a*, where it coincides with the original system (2.2). This condition will be verified *a posteriori* for the controlled trajectories considered below. Figure 2*b* shows two distinct solutions to the abseiling problem, one resulting from a simple feedback law and the other arising from a time-optimal control problem. While the time-optimal solution reaches *E* in finite time, the feedback-based solution only approaches *E* asymptotically as *t* → ∞.

These solutions will be studied in detail in the following sections. A trajectory of the uncontrolled system (*u* = 0), which does not solve the abseiling problem, is also shown for comparison.

### (b) Feedback control

In this section we solve the abseiling problem by constructing a state feedback controller *u* = *u*(***x***) for which every trajectory of the closed-loop system 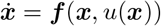 converges to the ray of fixed points *E*. The feedback controller should be biologically plausible, in that it should depend only on information that the spider can measure. We assume that this includes the silk strain *ε*, and therefore the excess strain *e*, but not the velocity or the unstretched silk length.

To motivate the feedback law, consider the energy-like function *V* (***x***) = (*le*^2^ + *v*^2^)*/*2, which is nonnegative and vanishes only on the ray of fixed points. In the taut region *e* ≥ −*a*, direct differentiation gives

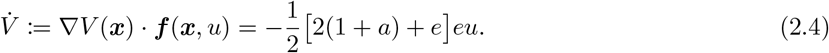

Thus *V* is conserved on taut intervals when *u* ≡0. Equation (2.4) suggests choosing *u* proportional to *e*, with the caveat that *u* must be nonnegative. We therefore consider the proportional state feedback law

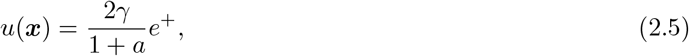

where *γ >* 0. While the dragline is taut, equation (2.4) then becomes

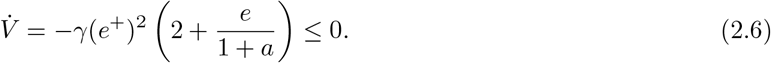

As shown in proposition 1 in appendix A(a), this feedback law successfully solves the abseiling task for every *γ >* 0: every trajectory of the closed-loop system converges to some point (0, 0, *l*_∞_) on the ray of fixed points. Figure 3*a*,*b* shows numerically computed trajectories of the full nonlinear system (2.2) under the proportional feedback law (2.5). The axes are normalized by the initial velocity *v*_0_ to reveal the weak dependence of the rescaled trajectories on *v*_0_: trajectories with the same value of *γ* but varying *v*_0_ approximately collapse onto a single curve under this change of variables. For smaller values of *γ*, however, the spider may experience a large maximum excess strain, especially for large *v*_0_. Increasing *γ* reduces the maximum excess strain and suppresses oscillations, but slows the asymptotic convergence and requires a larger total silk payout. This is made precise in proposition 2, which shows that there exists a critical gain *γ*_*c*_ = *γ*_*c*_(*v*_0_, *a*) such that, whenever *γ* ≥ *γ*_*c*_, the bound 0 ≤ *e*(*t*) ≤ *v*(*t*)*/γ* holds for all *t* ≥ 0. In this regime, the total silk payout is

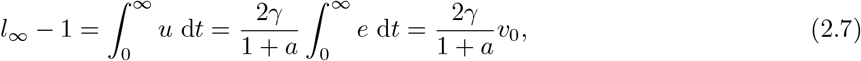

where the last equality follows from 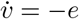 and *v*(*t*) →0. Moreover, since *v*(*t*) ≤*v*_0_, the maximum excess strain *e*_max_ := max_*t*_≥_0_ *e*(*t*) satisfies *e*_max_ ≤*v*_0_*/γ*. The trade-off between *e*_max_ and the total silk payout *l*_∞_− 1 is shown in figure 3*c*.

**Figure 3:**
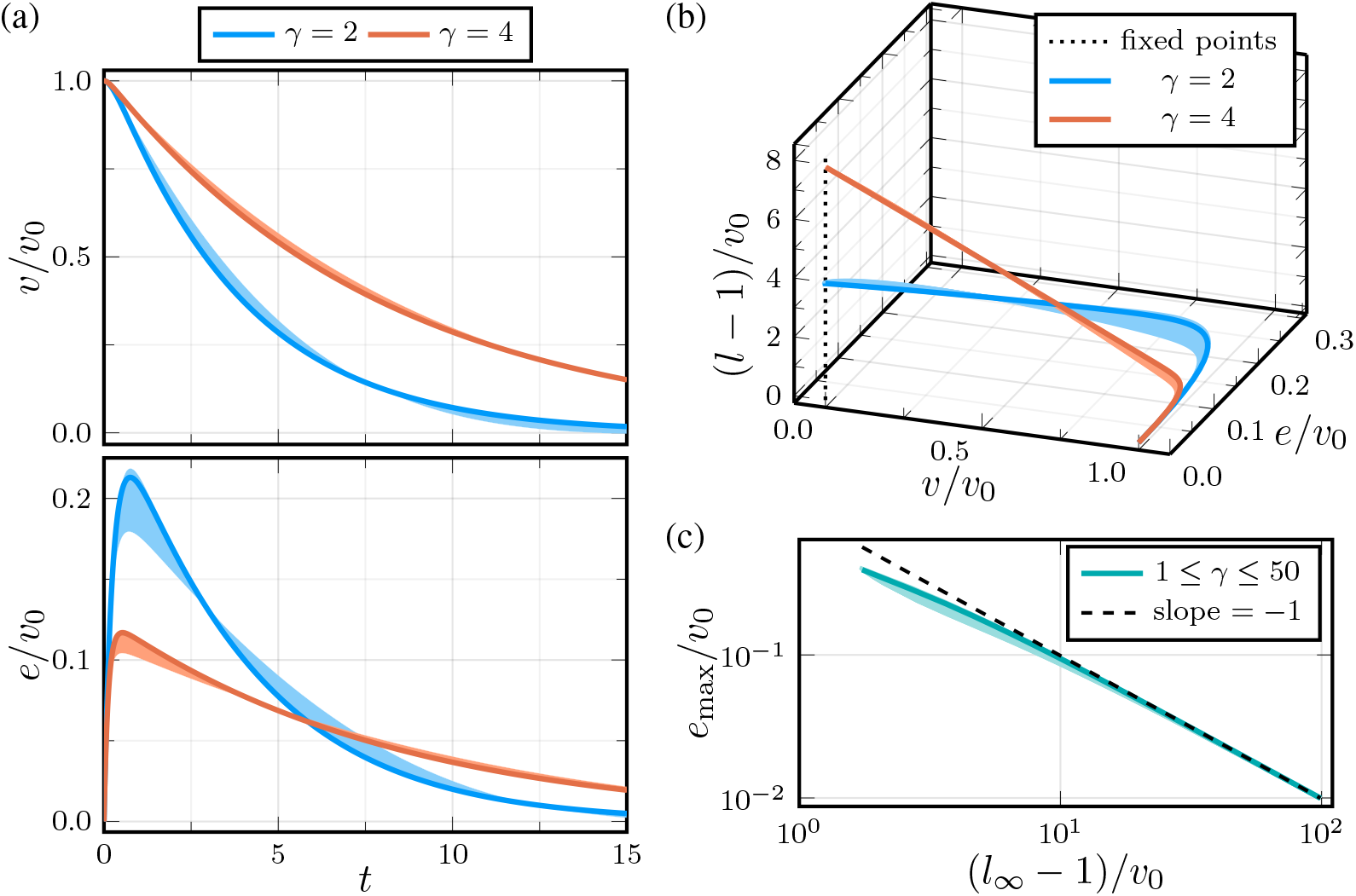
Trajectories of the nondimensional abseiling system (2.2) under the proportional feedback law (2.5). (a) Normalized velocity *v/v*_0_ and excess strain *e/v*_0_ as functions of nondimensional time *t* for *γ* = 2 and *γ* = 4. The solid curves correspond to 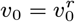. The shaded regions show the envelopes obtained by varying *v*_0_ over 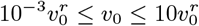. The same trajectories shown in the normalized state-space coordinates *v/v*_0_, *e/v*_0_, and (*l* 1)*/v*_0_, displaying convergence to the ray of fixed points *v* = *e* = 0, shown as a black dotted line. (c) The normalized maximum excess strain *e*_max_*/v*_0_ plotted against the normalized silk payout (*l*_*∞*_ *−* 1)*/v*_0_ for 1 *≤ γ ≤* 50. The solid curve corresponds to 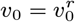, and the shaded region is obtained by varying *v*_0_ over 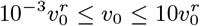. The dashed black line has slope − 1. Here *v*_0_ = *v*(0). In all panels, *a* = *g/K* = 0.005, and 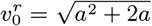. corresponds to a fall starting from rest at *y*=0.

### (C) Time-optimal control

#### Statement of the optimal control problem

The state feedback controller found in the previous section is inadequate in that the spider only comes to a stop asymptotically as *t*→∞. Moreover, as previously noted, the convergence is slower for larger values of *γ*, corresponding to smaller maximum strains. Therefore, this strategy is impractical if the breaking strain of the silk is small. Let us then pose the question of which control *u* accomplishes the abseiling task in minimum time, subject to the bounds *e*≤ *e*_*m*_ on the excess strain and *u*≤ *u*_*m*_ on the spinning rate. Let ***x***_0_ = (*v*_0_, 0, 1) be the initial state as before. For the analysis below, we use the simplified system (2.3). Although this system agrees with the original system (2.2) only in the taut region *e* ≥ − *a*, we verify below that the trajectories obtained in the parameter regimes of interest remain in this region. For a measurable control *u* : [0, ∞) → [0, *u*_*m*_], define the terminal time *t*_*f*_ = *t*_*f*_ (*u*) as the first time that the corresponding state trajectory ***x*** hits the ray of fixed points *E*. We say that *u* is admissible if *t*_*f*_ *<* ∞and the trajectory satisfies the inequality constraint

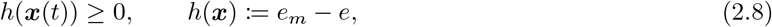

for 0 ≤ *t* ≤ *t*_*f*_. The abseiling time-optimal control problem (TOCP) is to find an admissible control *u* which minimizes *t*_*f*_. We denote the optimal control by *u*^*∗*^, the corresponding trajectory by ***x***^*∗*^, and the optimal terminal time by 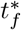.

Before applying the Pontryagin maximum principle, we make a few observations. If *e*_*m*_ and *u*_*m*_ are too small for a given initial velocity *v*_0_, then no control satisfies the strain constraint *e* ≤*e*_*m*_ and the problem is infeasible. Conversely, appendix A(b) shows that if the strain constraint is satisfied for the constant control *u* = *u*_*m*_, then an admissible control exists. The boundary of the infeasible region is therefore determined by the maximum strain attained under the constant control *u* = *u*_*m*_; in the small-*v*_0_ limit this is

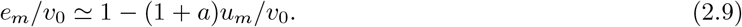

This region of parameter space is shown in gray in figure 4*b*.

**Figure 4:**
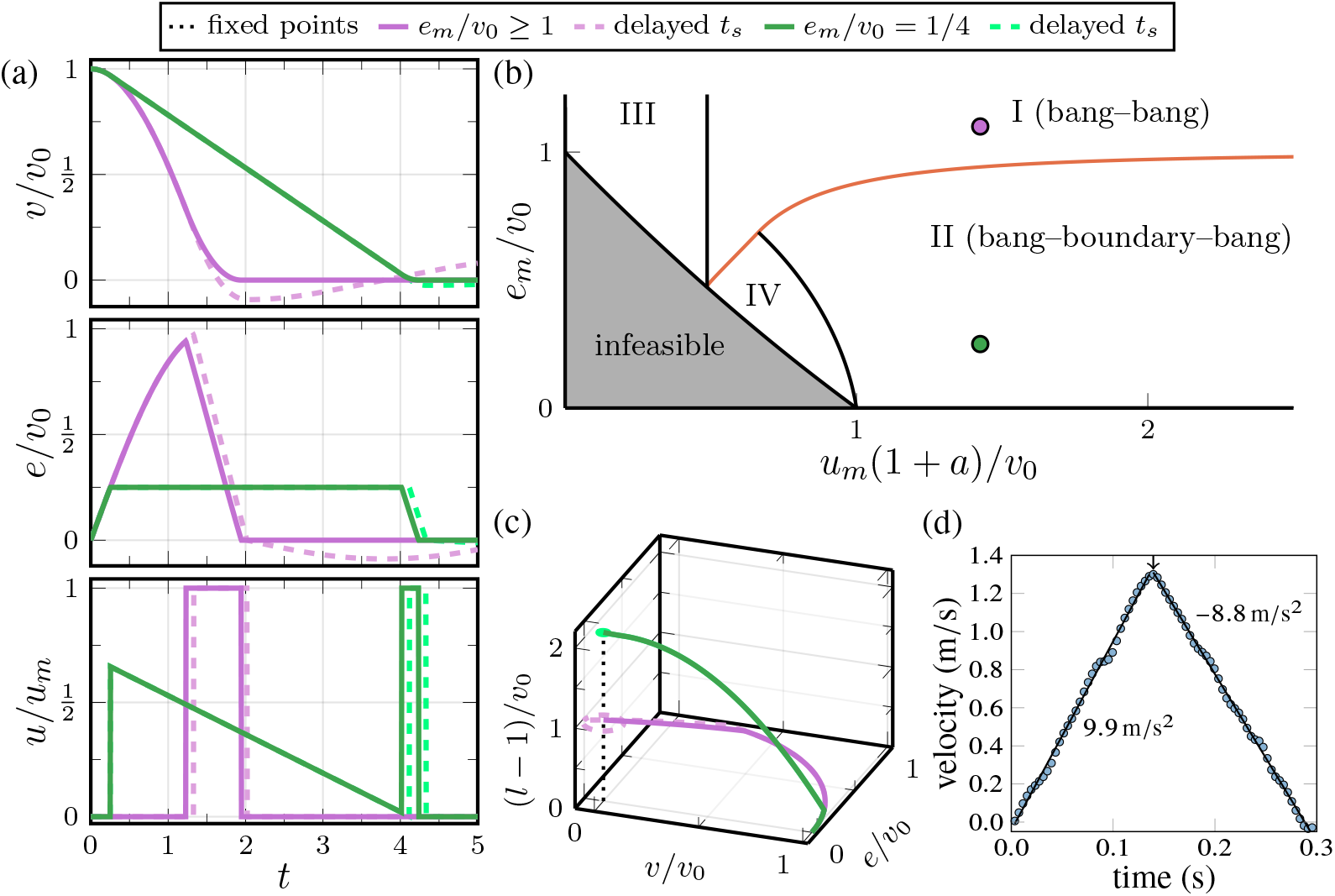
Time-optimal control of the nondimensional abseiling system (2.2) subject to the strain constraint *e*≤ *e*_*m*_ and spinning rate constraint *u* ≤*u*_*m*_. (a) Normalized velocity *v/v*_0_, excess strain *e/v*_0_, and spinning rate *u/u*_*m*_ as functions of nondimensional time *t*. For *e*_*m*_*/v*_0_ ≥1 (purple), the strain constraint is inactive and the numerically minimizing control is bang–bang. For *e*_*m*_*/v*_0_ = 1*/*4 (green), the trajectory includes a boundary arc with constant deceleration, 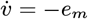. The dashed curves are obtained by delaying the switching time *t*_*s*_ by 0.1 units, showing that the trajectories pass near the ray of fixed points before undergoing small-amplitude oscillations. (b) Parameter space for the time-optimal control problem. The orange curve separates regions in which the strain constraint remains inactive along the minimizing trajectory from those in which the trajectory contains a boundary arc with *e* = *e*_*m*_. In regions I and II, numerical optimization gives the controls in equations (2.10) and (2.12), respectively. The purple and green points indicate the parameter values used in panels (a) and (c). In regions III and IV, the minimizing control involves additional bang and boundary arcs. In the gray region, the problem is infeasible. (c) The trajectories from panel (a) shown in the normalized state-space coordinates *v/v*_0_, *e/v*_0_, and (*l* −1)*/v*_0_. The ray of fixed points *v* = *e* = 0 is shown as a black dotted line. See also Supplementary Video S1. In panels (a)–(c), *a* = *g/K* = 0.01, *u*_*m*_ = 0.2, and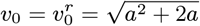, corresponding to a fall starting from rest. (d) Downward velocity of an *Araneus diadematus* during a vertical fall followed by abseiling. The fall was recorded at 250 frames/s. After a 0.13 s free fall, the spider engages its internal brake and decelerates at an approximately constant rate of 8.8 m/s^2^, consistent with the boundary arc in panel (a). The acceleration was estimated by least squares regression (solid lines). Data were digitized from Ortlepp and Gosline [7].

At the other extreme, if *e*_*m*_ ≥*v*_0_, the strain constraint does not restrict the relevant trajectories, and the problem is feasible for any *u*_*m*_ *>* 0. When *u*_*m*_ is sufficiently large, the system can be controlled to *E* by a one-switch bang–bang control of the form

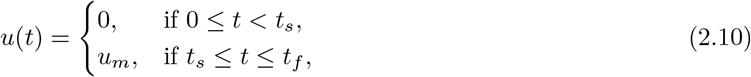

where *t*_*s*_ is a switching time to be determined.

For smaller *e*_*m*_, the bang–bang control (2.10) is not admissible because the trajectory exceeds the maximum allowed strain. For *e*_*m*_*/v*_0_ *<* 1, the initial *u* = 0 arc reaches *e* = *e*_*m*_ at time

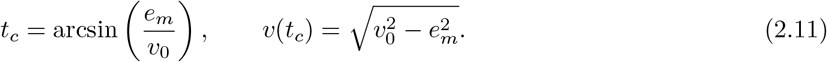

Since 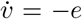, maintaining *e* = *e*_*m*_ thereafter allows the spider to continue decelerating at the maximum rate permitted by the strain constraint. The condition ė = 0 requires *u* = *v/*(1 + *a* + *e*_*m*_), leading to the bang–boundary–bang control

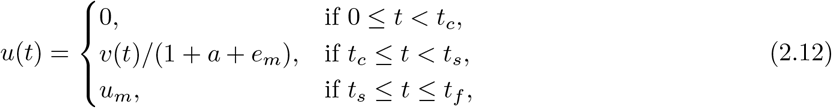

where *t*_*s*_ is a switching time to be determined. The spinning rate required to keep the strain fixed at *e*_*m*_ is largest when the constraint is first reached. The spinning-rate constraint is then satisfied precisely when

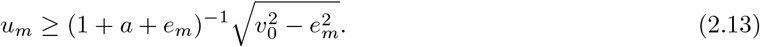

The controls (2.10) and (2.12) and their corresponding trajectories are shown in figure 4*a*,*c*. As we show below, they are the time-optimal controls over the parameter ranges of primary interest. The trajectories satisfy *e* ≥ 0 by construction, justifying the use of the simplified system (2.3). For the larger value of *e*_*m*_, shown in purple, the one-switch bang–bang control reaches the ray of fixed points without encountering the strain constraint. For the smaller value of *e*_*m*_, shown in green, the trajectory reaches *e* = *e*_*m*_ and maintains this value for a finite interval before switching to *u* = *u*_*m*_. The latter trajectory maintains a substantially smaller maximum strain, at the cost of a longer terminal time.

#### Pontryagin maximum principle

The Pontryagin maximum principle (PMP) provides necessary conditions for a control-trajectory pair to solve the TOCP. We use a formulation for problems with pure state constraints [13]; details are given in appendix A(b). The PMP associates with an optimal trajectory a costate trajectory ***λ*** = (*λ*_*v*_, *λ*_*e*_, *λ*_*l*_) and a constant cost multiplier *λ*_0_ ≥ 0. The optimal control is determined by maximizing the Hamiltonian *H*(***x***, *u, λ*_0_, ***λ***) = ***λ*** · (***F*** (***x***) + ***G***(***x***)*u*) −*λ*_0_ over the admissible values of *u*, taking the strain constraint into account. In our case, the Hamiltonian depends on *u* only through the *switching function*

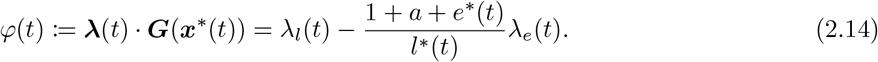

On an *interior arc*, i.e. an interval on which the strain constraint is not active, the PMP requires *u*^*∗*^(*t*) = *u*_*m*_ when *φ*(*t*) *>* 0 and *u*^*∗*^(*t*) = 0 when *φ*(*t*) *<* 0. Thus *u*^*∗*^ switches between its minimum and maximum values at switching times when *φ* changes sign. An interior arc on which *φ* vanishes identically is called a *singular arc*; appendix A(b)(ii) shows that such arcs do not occur for this system. An interval on which the strain constraint is active is called a *boundary arc*. On such an arc, *e* = *e*_*m*_, and maintaining the strain at this value requires *ė* = 0. It follows from equation (2.3) that *u*^*∗*^(*t*) = *v*^*∗*^(*t*)*/*(1 + *a* + *e*_*m*_). Thus, the PMP implies that an optimal control consists only of interior arcs where the control is bang–bang and boundary arcs that maintain *e* = *e*_*m*_. Moreover, it can be shown that the final arc must have *u*^*∗*^ = *u*_*m*_. This determines the optimal control up to the number and ordering of the arcs, as well as the switching times.

The full PMP conditions yield a multi-point boundary value problem which could in principle be solved to determine the optimal control [13]. Here, we instead use a simpler approach based on optimization over the switching times [14]. Using this approach, we obtain the parameter-space diagram shown in figure 4*b*. In region I, the time-optimal control is the one-switch bang–bang control (2.10), while in region II it is the bang–boundary–bang control (2.12). In regions III and IV, these simple control structures are no longer sufficient to reach, and additional bang and/or boundary arcs are required. We do not analyze these regimes further.

The boundaries in figure 4*b* mark the parameter values where the corresponding control structures no longer reach *E* without violating the constraints. At the boundary between regions I and III, the maximum spinning rate becomes too small for a one-switch bang–bang trajectory to reach *E*; in the small-*v*_0_ limit, this happens when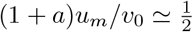. The remaining boundary of region I is determined by the maximum strain attained along the one-switch bang–bang trajectory. Finally, the boundary between regions II and IV occurs when the spinning rate required at the beginning of the boundary arc reaches *u*_*m*_. As shown in appendix A(b), this boundary is given exactly by equality in equation (2.13). The boundaries plotted in figure 4*b* are computed from the full nonlinear system (2.2). Although exact expressions are not available for all of the region boundaries, they are readily approximated in the small-*v*_0_ limit. This limit is relevant because *v*_0_ is small for the typical parameter values discussed in section 2(a), while numerical solutions show that the boundaries depend only weakly on *v*_0_ over the relevant range. The resulting asymptotic expressions are derived in appendix A(b).

The feedback and time-optimal controls offer different ways of accomplishing the abseiling task. The feedback law of section 2(b) is robust to perturbations, but the spider approaches the ray of fixed points only asymptotically, with smaller maximum strains requiring slower convergence and greater silk payout. In contrast, the time-optimal controls bring the spider to rest in finite time. The bang–bang control in region I does so rapidly, but requires a relatively large maximum strain and is sensitive to errors in the switching time. As illustrated by the dashed purple curves in figure 4*a*,*c*, slightly delaying *t*_*s*_ causes the trajectory to miss *E* and undergo oscillations. The bang–boundary–bang control in region II maintains the prescribed maximum strain at the cost of a longer terminal time, and is more robust to perturbations in *t*_*s*_. Supplementary Video S1 compares the feedback and time-optimal trajectories. Finally, we note that during the boundary arc the spider decelerates at a constant rate 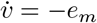. This behavior is consistent with the approximately constant deceleration observed by Ortlepp and Gosline [7], shown in figure 4*d*. In the physically realistic case, *e*_*m*_ is moderate or small and *u*_*m*_ is of order *v*_0_ or larger, placing the problem in region II.

## 3. The jumping task

### (a) Mathematical model

A minimal model of a jumping spider during the aerial phase is shown in figure 5. We assume that the motion is restricted to the *xy*-plane, where the *y*-axis points vertically upward, and approximate the spider’s body as an axisymmetric rigid body of mass *m*. The spider’s center of mass 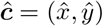 and the position of the spinnerets ŝ both lie on the body axis and are separated by a distance *d*. We denote by *θ* the angle that the body axis makes with the horizontal, so that ***ŝ*** = ***ĉ*** *d****e***_*θ*_, where ***e***_*θ*_ := (cos *θ*, sin *θ*). The spinnerets are connected to the origin by a dragline with the same constitutive law as in the abseiling model. Let 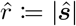 denote the distance from the origin to the spinnerets and *ϕ* the angle that the dragline makes with the horizontal. We define 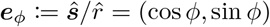 as the unit vector pointing from the origin toward the spinnerets. As in the abseiling section, we denote the unstretched silk length by 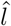 and the silk spinning rate by 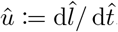. Neglecting air resistance, the spider experiences a gravitational force ***F***_*g*_ = *mg****e***_*y*_ and a silk force directed toward the origin:

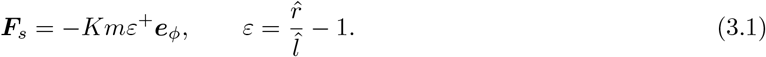

**Figure 5:**
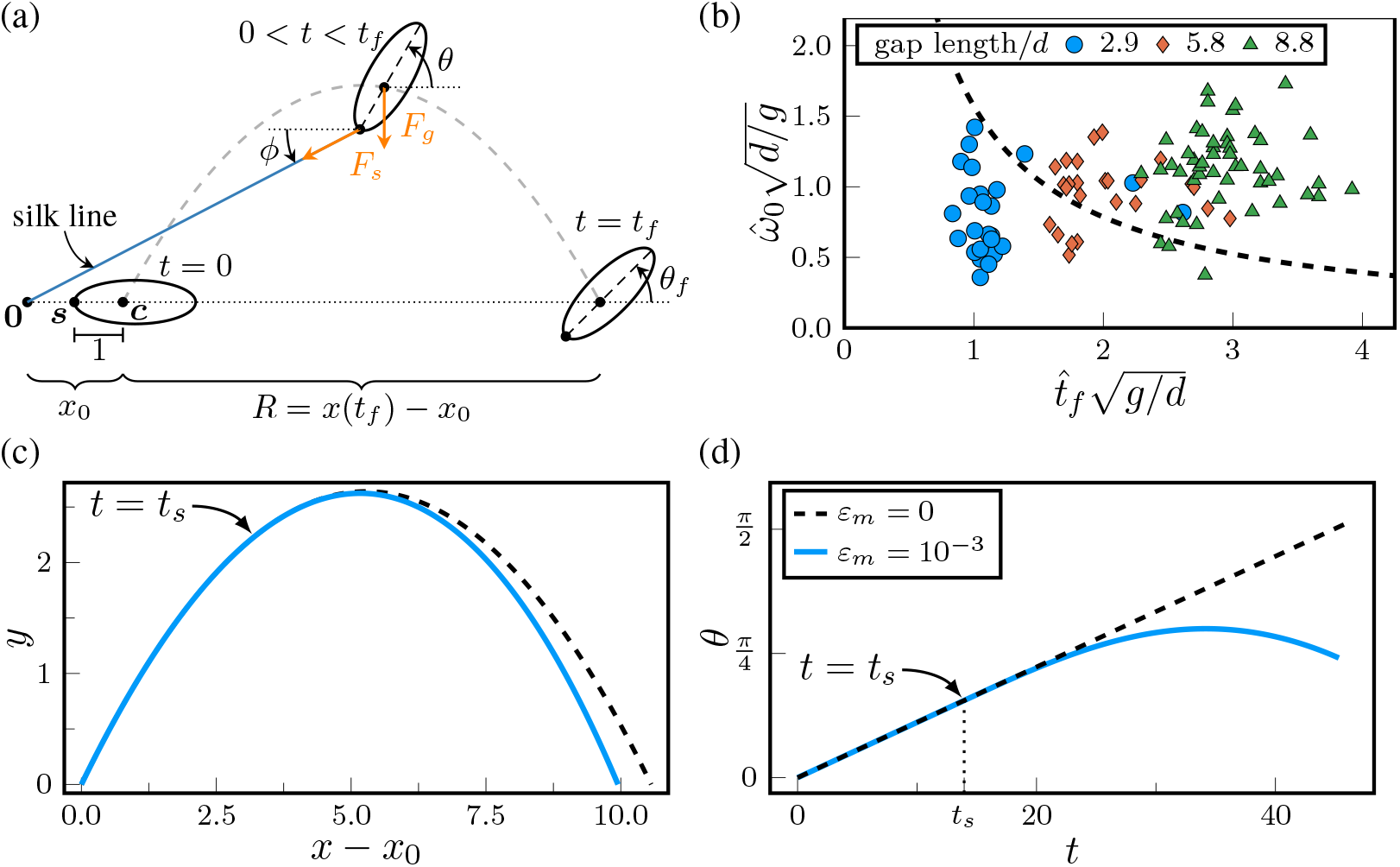
(a) Schematic of the jumping task in nondimensional variables. The spider is modeled as an axisymmetric rigid body of mass *m*, with its motion restricted to a vertical plane. The spinneret position ***s*** and center of mass ***c*** lie on the body axis and are separated by unit distance. The spinnerets are connected to the origin by a silk dragline, which makes an angle *ϕ* with the horizontal. The arrows indicate the silk and gravitational forces. The spider is shown at takeoff (*t* = 0), during the aerial phase (0 *< t < t*_*f*_), and at landing (*t* = *t*_*f*_). The body axis makes an angle *θ* with the horizontal, is the landing angle, and *R*= *x*(*t*_*f*_) − *x*_0_ is the horizontal range. (b) Nondimensionalized flight times 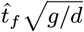 and initial angular velocities 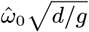 for *S. scenicus* jumping trials with different gap lengths. For an initially horizontal body, trials below the curve 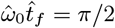 (dashed black) do not require a braking torque from the silk to avoid rotating beyond *θ*_*f*_ = *π/*2. (c) Simulated trajectories without silk braking (*ε*_*m*_ = 0, dashed black) and with silk braking (*ε*_*m*_ = 10^*−*3^, solid blue). See also Supplementary Video S2. (d) Body angle *θ* as a function of time for the same trajectories. The arrows in (c)–(d) mark the switching time *t* = *t*_*s*_, chosen as the first time at which positive silk tension produces a braking torque. In (c)–(d), *a* = 0.01, *j* = 0.25, *x*_0_ = 1, *ω*_0_ = 0.035, and *v*_0_ = 0.325, with launch angle *π/*4.

Here, as before, *ε* denotes the silk strain. As in the abseiling model, we require *û* = 0 when *ε <* 0, although this constraint is inactive for the trajectories considered here. We write the moment of inertia as *J* = *jmd*^2^, where *j* is a dimensionless coefficient that may in general depend on the spider’s posture; we approximate *j* as constant.

We nondimensionalize distances by *d* and time by 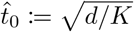:

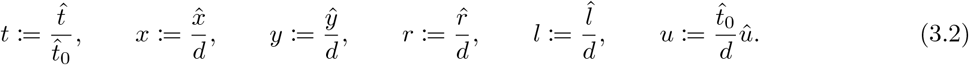

Force and moment balance then give the nondimensional equations of motion for the state variables 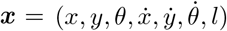:

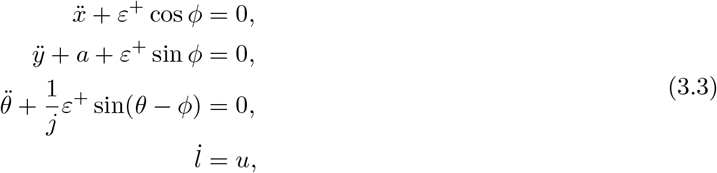

where *a* := *g/K* and dots denote derivatives with respect to the nondimensional time *t*. For simplicity, we consider only initial conditions with *y*(0) = 0, *θ*(0) = 0, and *l*(0) = *x*_0_ − 1, where *x*_0_ := *x*(0) ≥1 is left free.^1^ The condition on *l*(0) corresponds to a dragline that is taut and unstrained at takeoff. We denote the takeoff speed and initial angular velocity by *v*_0_ and *ω*_0_, respectively, and restrict attention to *ω*_0_ *>* 0. We define the landing time *t*_*f*_ as the first time *t >* 0 at which the center of mass returns to *y* = 0. The *jumping task* is to choose the spinning rate *u*(*t*) such that the terminal angle *θ*_*f*_ := *θ*(*t*_*f*_) satisfies 0 ≤*θ*_*f*_ ≤*π/*2, so that the spider lands upright. Without silk braking (*ε* ≡0), the trajectory is ballistic and 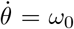 is constant. Assuming a launch angle of *π/*4, the total body rotation is

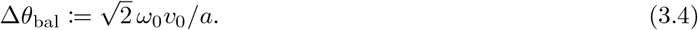

Thus, in the absence of other torques, silk braking is required precisely when Δ*θ*_bal_ exceeds *π/*2.

To compare this ballistic criterion with observed jumping kinematics, we analyzed high-speed recordings (2500 frames/s) of 100 jumps by 10 *S. scenicus* individuals across gaps of 1, 2, and 3 cm. Figure 5*b* compares the measured flight times 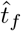 and initial angular velocities 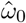, with the gap lengths expressed in units of *d*.

For this comparison, *d* is estimated by the mean spinneret–pedicel distance across all trials. For an initially horizontal spider, the dashed curve 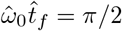 marks the boundary above which a braking torque is required to land upright. A simple estimate of aerodynamic damping suggests that air resistance reduces the total rotation by roughly 5–15% for representative parameters (appendix B(a)). Thus, air resistance alone cannot account for an upright landing for trials lying appreciably above the dashed boundary. This is consistent with previous evidence that torque from the dragline plays a role in controlling body orientation during flight [8,9].

### (b) Constant-strain control

Motivated by these observations and by the form of the time-optimal control for the abseiling task, we ask whether the spider can successfully land upright using a control which allows the dragline strain to increase to a target value *ε*_*m*_ after some switching time *t*_*s*_, and then keeps it constant:

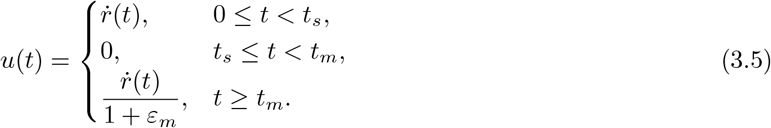

Here *t*_*m*_ denotes the first time *t*≥ *t*_*s*_ at which the strain reaches the target value *ε*_*m*_ ≥0. Initially, the spider freely lets out silk so that the dragline remains unstrained. After the switching time, no silk is spun until the strain reaches the target value *ε*_*m*_, after which spinning resumes at the rate required to maintain constant strain. A natural choice for *t*_*s*_, which we adopt throughout, is the moment when sin(*θ*− *ϕ*) becomes positive, so that positive silk tension produces a braking torque:

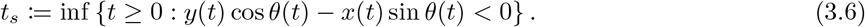

To demonstrate that this strategy can successfully complete the jumping task, figures 5*c*,*d* compare trajectories with and without silk braking for an initial condition chosen so that Δ*θ*_bal_ *> π/*2. The two trajectories coincide up to the switching time *t*_*s*_. Once the silk brake is activated, the resulting torque reduces the spider’s angular velocity and allows it to land upright. Supplementary Video S2 shows the corresponding jumping dynamics, highlighting how dragline tension generates a braking torque that limits body rotation and promotes an upright landing.

To assess the effectiveness of this strategy for stabilizing the spider when jumping across large gaps, we ask what target strain is required for a given initial condition. Let 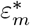 denote the minimum value of *ε*_*m*_ for which the control (3.5) satisfies the landing-angle constraint. Numerical simulations of equation (3.3) give the minimum-strain surface shown in figure 6*a*, with 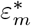 plotted as a function of the takeoff speed *v*_0_ and initial angular velocity *ω*_0_. Intuitively, the required strain increases with angular velocity, whereas its dependence on takeoff speed is non-monotonic. At fixed *ω*_0_, increasing *v*_0_ increases the flight time as *t*_*f*_ ~ *v*_0_. The rotation due to the initial angular velocity then scales as *ω*_0_*t*_*f*_ ~ *ω*_0_*v*_0_, whereas the angular correction produced by the silk torque scales approximately as 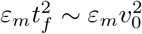. The competition between these two scalings gives rise to the non-monotonic dependence on *v*_0_. To characterize the required strain in the physically relevant region near the zero-strain boundary, we transform to the coordinates

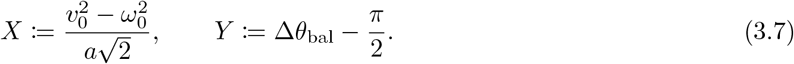

**Figure 6:**
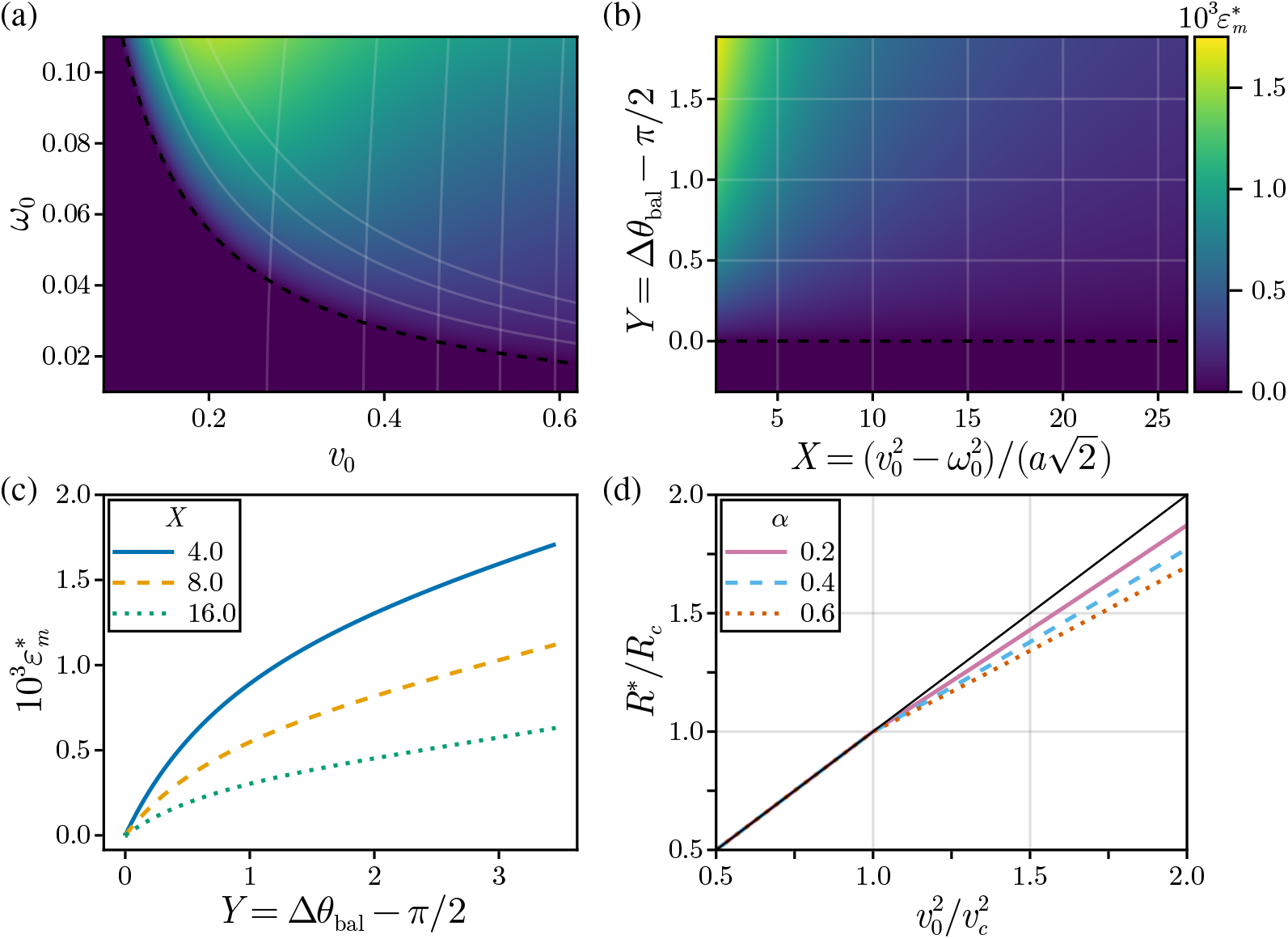
Minimum silk strain required to satisfy the landing-angle constraint 0 *≤θ*_*f*_ *≤π/*2 and its effect on horizontal range *R*. (a) Minimum target strain 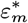 as a function of the takeoff speed *v*_0_ and initial angular velocity *ω*_0_. The total body rotation along the corresponding ballistic trajectory is 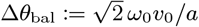. The dashed black curve marks the zero-strain boundary Δ*θ*_bal_ = *π/*2; below this curve, the landing-angle constraint is satisfied without silk torque. (b) The same minimum-strain surface in the transformed coordinates 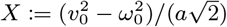 and *Y* := Δ*θ*_bal_ *− π/*2. The grid in (a)–(b) shows corresponding curves of constant *X* and constant *Y*. (c) Minimum target strain 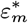 as a function of *Y*, for fixed values of *X*. (d) Horizontal range *R*^*∗*^ of a jump with minimum silk strain 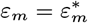. The scaled horizontal range *R*^*∗*^*/R*_*c*_ is shown as a function of 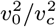, where *v*_0_ and *ω*_0_ are varied simultaneously with *α* = *ω*_0_*/v*_0_ held constant. Here, for each fixed, *α v*_*c*_ is the takeoff speed corresponding to Δ*θ*_bal_ =*π/*2, and 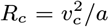 is the corresponding horizontal range. The black line shows the range of a ballistic trajectory with the same *v*_0_ and *ω*_0_. In all panels, *a* = 0.01, *j* = 0.25, *x*_0_ = 1, and the launch angle is *π/*4.

Up to a uniform scaling and translation, this transformation is the conformal map *z* ↦*z*^2^ (where *z* = *v*_0_ +*iω*_0_), and therefore provides natural orthogonal coordinates with the zero-strain boundary mapped to *Y* = 0 (figure 6*b*). In these coordinates, the minimum target strain is monotonic in each variable. Figure 6*c* shows the dependence of 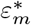 on *Y* at fixed *X*.

We finally ask whether this strategy leads to greater horizontal range, or whether the braking required to control body rotation offsets the gain from a higher takeoff speed. The data in figure 5 show substantial variation in initial angular velocity, but suggest that *ω*_0_ may increase with *v*_0_, particularly for longer jumps. To account for this possibility, we consider a family of initial conditions *ω*_0_ = *αv*_0_, with *α >* 0. Under this relation, there is a maximum takeoff speed *v*_*c*_ for a jump without silk braking and a corresponding maximum range *R*_*c*_, given by

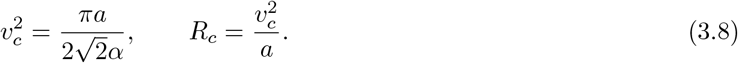

For fixed *α*, we vary *v*_0_ (and hence *ω*_0_) and compute the range *R*^*∗*^ of a jump using the minimum target strain 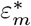. Numerical simulations in figure 6*d* show that, above *v*_*c*_, the range *R*^*∗*^ continues to increase with *v*_0_ and is only slightly reduced relative to that of a ballistic trajectory with the same takeoff speed. In appendix B(b), we show analytically that the range can be increased beyond *R*_*c*_ for a range of *α* relevant to the observed jumping behavior. Thus, the silk torque allows the spider to exploit higher takeoff speeds for greater range while landing upright.

## 4. Discussion

The use of dragline silk to regulate locomotion poses an unusual control problem: the spider steers its trajectory by dynamically deploying a mechanical element, via a particularly simple form of embodied actuation, sensing, and control. Here we have considered two manifestations of this problem—control of descent during abseiling and of body orientation during jumping—and shown that both can be framed in terms of regulating a common mechanical quantity, the dragline’s strain. In both cases, the spider can rely on feedback based on the strain alone, without requiring an estimate of its translational or angular velocity. The dragline can therefore be viewed not just as a tether, but as part of a dynamical sensorimotor loop coupling the spider’s body, the spinnerets, and the environment.

For abseiling, the analysis reveals two complementary control strategies. A proportional feedback law based on the silk strain brings the spider to rest asymptotically, with a trade-off between the maximum strain and the convergence rate. The time-optimal formulation makes this trade-off explicit by minimizing the stopping time subject to an upper bound on the strain. The nearly constant deceleration observed experimentally then arises naturally from the constant-strain phase of the time-optimal control. The presence of mechanoreceptors on the spinnerets that are sensitive to dragline tension suggests that such a strategy could be implemented through a simple strain-regulating reflex. The same principle extends to the jumping task, where maintaining a small target strain can prevent over-rotation with only a modest reduction in horizontal range. These results raise the possibility that the in-flight braking response of salticids evolved as a specialization of a more general strain-regulation response used during descent. This adaptation could allow jumping spiders to exploit higher takeoff speeds for greater range while maintaining control of their landing orientation.

The simplicity of our models also defines their limitations. Incorporating rate-dependent silk mechanics, active postural changes, and aerodynamic forces would help disentangle the relative contributions of passive mechanics, active body motion, and silk regulation. Accounting for delays and noise in sensory feedback would further assess the robustness and biological plausibility of the proposed control strategies. Comparative experiments during abseiling and jumping could test whether silk release is regulated by similar strain-dependent feedback in the two behaviors, while measurements or perturbations of the underlying sensory and muscular systems could indicate whether they share a common physiological basis. Measurements of dragline tension across trials with different initial conditions could provide a direct test of the strain-regulation hypothesis in both abseiling and jumping.

## Data accessibility

All data, simulation code, and Supplementary Videos S1 and S2 associated with this paper are available from Zenodo [15].

## Acknowledgements.

We thank the members of the Mahadevan lab for helpful discussions.

## A. Analysis of the abseiling task

We use the notation established in section 2. Recall that the abseiling spider is described by the system (2.2) which, in the taut region *e* ≥ − *a*, reduces to the control-affine system (2.3). For this reduced system, we will repeatedly use the energy-like function *V* (***x***) = (*le*^2^ + *v*^2^)*/*2 whose derivative is given by equation (2.4). Along a trajectory, we write *V* (*t*) := *V* (***x***(*t*)) for convenience.

### (a) Feedback control

Under the feedback law (2.5), the full closed-loop system is

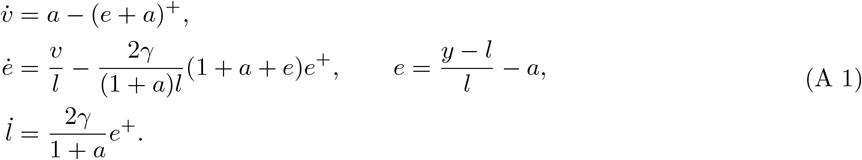

Unless otherwise stated, the initial condition is taken as (*v*(0), *e*(0), *l*(0)) = (*v*_0_, *e*_0_, 1), where *l*(0) = 1 is fixed by the nondimensionalization while *v*_0_, *e*_0_ are general. Depending on the initial condition and the value of *γ*, the trajectory may remain in the positive half-space *e*≥0 for all time, or it may alternate between *positive excursions* with *e >* 0, and negative excursions with *e <* 0. This distinction is useful because the control is active and energy is dissipated only on positive excursions, whereas the dynamics are conservative and *l* is constant on negative excursions. We denote the velocity and silk length at the start of the *n*th positive excursion by *v*^(*n*)^ and *l*^(*n*)^, respectively. Note that the same positive excursion ends with *v* = − *v*^(*n*+1)^, *l* = *l*^(*n*+1)^.

We now state the main convergence result:

#### Proposition 1

(Global convergence). *For every γ >* 0, *every solution of* (A 1) *with l*(0) *>* 0 *converges to some point* (0, 0, *l*_*∞*_) ∈ *E*, *where l*_*∞*_ *>* 0 *depends on γ and the initial condition*.

The result follows readily from LaSalle’s invariance principle if we can prove that *l* is bounded. To that end, we will use two lemmas. The first shows that each trajectory eventually remains in the taut region *e* ≥ −*a*, and records useful identities for the positive excursions.

#### Lemma 1

(Excursion identities and eventual tautness). *For every solution of* (A 1), *there exists T >* 0 *such that*

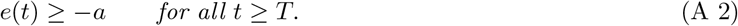

*Moreover, if the n-th positive excursion begins at e* = 0, *v* = *v*^(*n*)^ *>* 0, *l* = *l*^(*n*)^, *and ends at e* = 0, *v* = −*v*^(*n*+1)^, *l* = *l*^(*n*+1)^, *then*

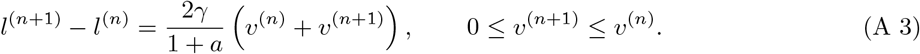

The second lemma is a technical estimate which bounds the increase in *l* over a single positive excursion.

#### Lemma 2

(Length increment bound for large *l*). *Fix a solution of* (A 1), *and consider its positive excursions. For all excursions n such that l*^(*n*)^ *is sufficiently large*,

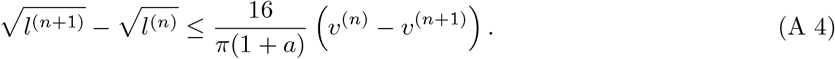

*Proof of proposition 1*. By (A 2), after some finite time *T*, the trajectory remains in the region *e* ≥ −*a*.

Therefore, from time *T* onward, the first equation of (A 1) reduces to 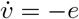 and the energy identity (2.6) holds, with 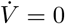 exactly when *e* ≤ 0.

We first observe that if *l*(*t*) is bounded, then the trajectory is precompact. Indeed, if *l*(*t*) ≤ *L* for *t* ≥ *T*, then 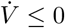 for *t* ≥ *T*, and hence the forward orbit {***x***(*t*) : *t* ≥ *T*} is contained in the compact set

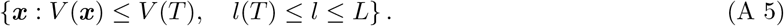

Applying LaSalle’s invariance principle [16, § 4.2] to the closure of the forward orbit, the trajectory approaches the largest invariant subset of 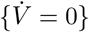. On 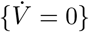 = {*e* ≤ 0}, *l* is constant, so *e* satisfies *ë*+ *e/l* = 0. Every nontrivial solution exits *e*≤0, so the largest invariant subset is contained in *E*. It remains only to show that *l*(*t*) is bounded.

Since 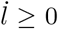, either *l*(*t*) converges to a finite limit *l*_*∞*_ or *l*(*t*) → +∞. First suppose that there are only finitely many positive excursions. Then, after some time 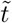, either the solution is already on *E*, or it remains in the half-space *e* ≥ 0 for all later times. In the latter case, for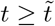,

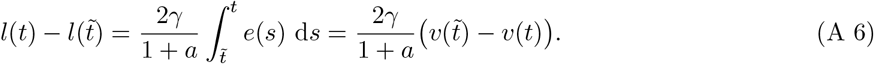

The right-hand side is bounded because *v*(*t*) is bounded by 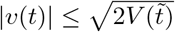. Hence *l*(*t*) is bounded.

Now suppose that there are infinitely many positive excursions. If *l*(*t*) were unbounded, then *l*^(*n*)^ → ∞.

Hence (A 4) applies for all sufficiently large *n*. Therefore, for all *M* ≥ *N*, with *N* sufficiently large,

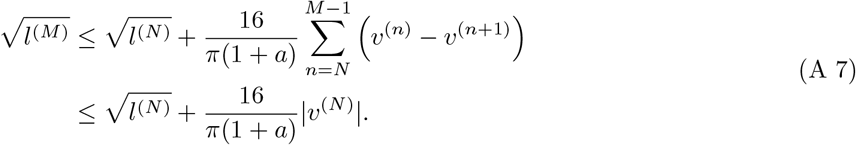

This contradicts *l*^(*M*)^ → ∞. Hence *l*(*t*) is bounded also in the infinite-excursion case, which concludes the proof. □

*Proof of lemma 1*. On a positive excursion, equation (A 1) gives

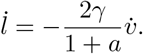

Integrating between the start and end of the positive excursion gives the length increment in (A 3). Similarly, integrating equation (2.6) for 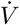 over the positive excursion gives

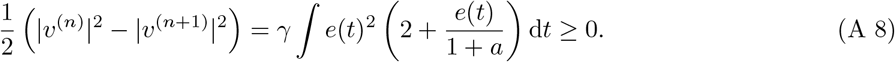

Hence *v*^(*n*+1)^ ≤ *v*^(*n*)^, completing the proof of (A 3).

It remains to prove that the trajectory eventually remains in the taut region *e*≥−*a*. If there are only finitely many positive excursions, then the result is clear: Since no trajectory can remain in *e <* 0 for all time, the solution must eventually remain in *e* ≥0. Now suppose instead that there are infinitely many positive excursions, and consider the negative excursion following the *n*-th positive excursion. While − *a* ≤*e* ≤ 0, the dynamics reduce to the harmonic oscillator 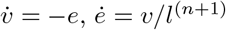, and a simple calculation shows that the slack region is reached if and only if

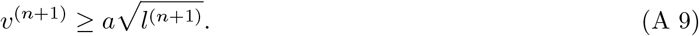

We now have two cases. If *l*^(*n*)^ → ∞, then the inequality fails for sufficiently large *n* since *v*^(*n*)^ ≤ *v*^(1)^. If instead *l*^(*n*)^ → *l*_*∞*_ is bounded, then summing over *n* in (A 3) gives

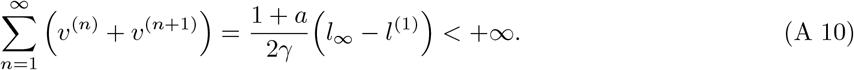

Therefore *v*^(*n*+1)^ →0. On the other hand, 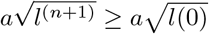, so (A 9) eventually fails. Thus in either case the trajectory enters *e <* −*a* only finitely many times. □

*Proof of lemma 2*. Consider a positive excursion with *l*^(*n*)^ ≥ *L*, where *L* will be chosen later. Let 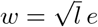 and introduce polar coordinates in the (*v, w*)-plane:

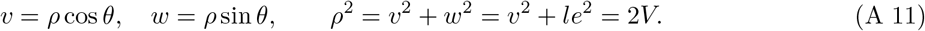

Since *ρ >* 0 throughout the excursion, the transformation is well-defined. We have *θ* = 0, *ρ* = *v*^(*n*)^ at the start of the excursion, and *θ* = *π, ρ* = *v*^(*n*+1)^ at the end. Our aim is to estimate the decrease *v*^(*n*)^ − *v*^(*n*+1)^ by obtaining a bound for d log *ρ/* d*θ* and integrating over the excursion. Indeed, if we can show that

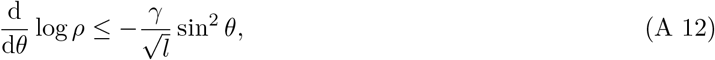

then integrating from *θ* = 0 to *θ* = *π* and using *l* ≤ *l*^(*n*+1^) yields

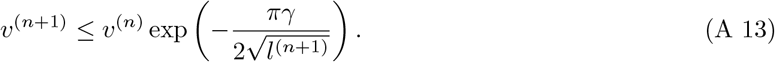

By increasing *L* if necessary, we can ensure that the exponent is at most 1. Setting 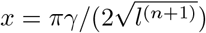and using 1 − *e*^*−x*^ ≥ *x/*2 for 0 ≤ *x* ≤ 1, we obtain

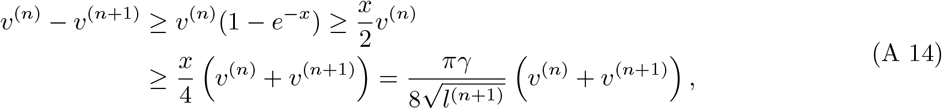

where we used *v*^(*n*+1)^ *v*^(*n*)^ in the third step.

Combining (A 14) with (A 3) gives

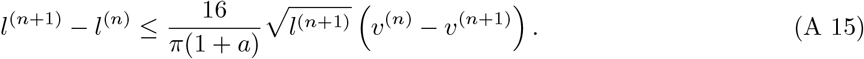

Dividing by 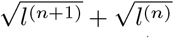 yields the result (A 4). Thus it remains to prove the bound (A 12).

First, from equations (A 11) and (2.6), and noting that *a >* 0, we have

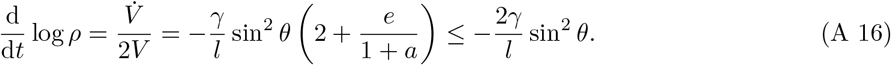

To obtain the desired bound, it is enough to show that

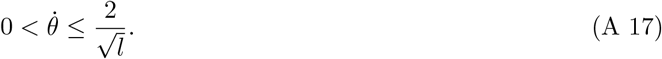

Differentiating and using equation (A 1), we find

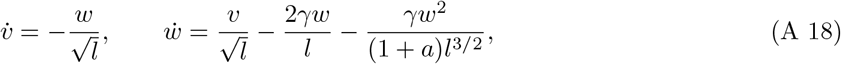

and

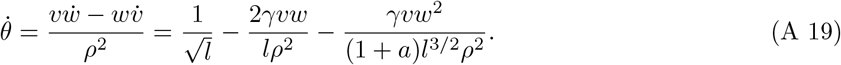

Rearranging and using 2|*vw*| ≤ *ρ*^2^, |*v*|*w*^2^*/ρ*^2^ ≤ *ρ* ≤ *v*^(1)^, we find

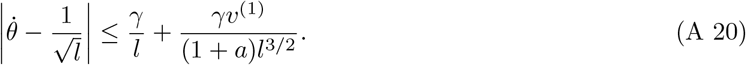

If we choose *L* sufficiently large, every positive excursion with *l*^(*n*)^ ≥ *L* therefore satisfies (A 17) throughout the excursion. Combining with (A 16) gives the desired bound

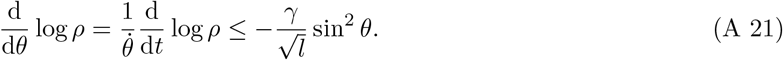

The next result shows that if *γ* is chosen sufficiently large, then a solution starting from the standard initial condition ***x***(0) = (*v*_0_, 0, 1) remains in the half-space *e* ≥ 0 for all time.

#### Proposition 2

(No overshoot for sufficiently large *γ*). *Assume e*(0) = 0, *l*(0) = 1, *and v*(0) = *v*_0_ *>* 0. *Define*

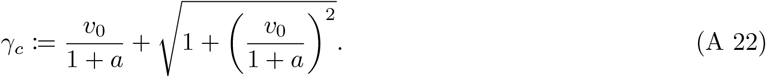

*If γ* ≥ *γ*_*c*_, *then the trajectory remains in the cone*

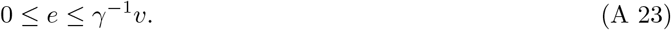

*In particular, e*(*t*) ≥ 0 *for all t* ≥ 0, *and it follows that*

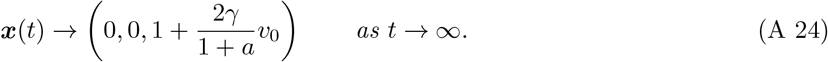

*Proof*. The initial condition satisfies *e*(0) = 0, so it lies in the cone (A 23). It suffices to show that the trajectory cannot exit the cone through either of the boundaries *e* = 0 or *e* = *γ*^*−*1^*v*. For the lower boundary, *e* = 0, this is immediate since there *ė* = *v/l* ≥ 0 and thus the vector field points inward everywhere. On the upper boundary, *e* = *γ*^*−*1^*v*, a short calculation gives

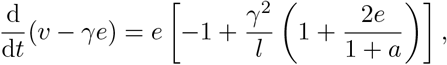

and thus the vector field points inward whenever *l ≤ γ*^2^.

Now suppose that the trajectory has a first exit through the upper boundary. Up to this first exit, we have *e* ≥ 0 and equations (A 1) yield the conserved quantity

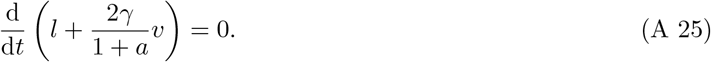

Thus (using *v* ≥ 0),

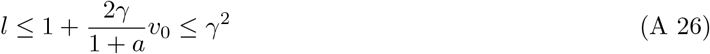

where the second inequality follows from the assumption *γ*≥ *γ*_*c*_. Therefore *l* ≤ *γ*^2^, so the vector field points inwards where the trajectory exits through the upper boundary, a contradiction.

This shows that the trajectory is contained in the cone (A 23), and in particular *e*(*t*) ≥ 0 for all *t* ≥ 0. By proposition 1, *v*(*t*) → 0 and *e*(*t*) → 0. Using the conserved quantity (A 25) and passing to the limit *t* → +∞ yields the second claim,

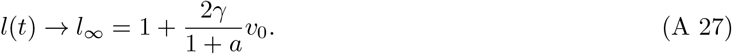

*Remark*. Linearizing the closed-loop dynamics around a point (0, 0, *l*) on the ray of fixed points E, from the side *e* ≥ 0, gives a zero eigenvalue corresponding to motion along E, and two nonzero eigenvalues

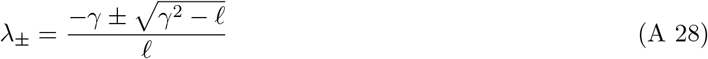

corresponding to motion in the (*v, e*)-directions. Thus the linearized dynamics are non-oscillatory when *γ*^2^ ≥*l*. Now assume we are in the no-overshoot regime and linearize around the limiting fixed point by choosing *l* = *l*_∞_. Then, the non-oscillation condition *l*_∞_≤ *γ*^2^ is equivalent to *γ* ≥ *γ*_*c*_ which is precisely the sufficient condition in proposition 2.

### (b)Time-optimal control

We consider the time-optimal control problem introduced in section 2(c), with initial condition ***x***(0) = (*v*_0_, 0, 1) and target set *E*, and write *U* = [0, *u*_*m*_] for the control set. As in the main text, we use the simplified system (2.3) for the analysis below.

#### (i)PMP conditions and arc structure

We use the control-affine representation 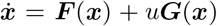, with ***F*** and ***G*** as defined in equation (2.3).

To account for the pure state constraint *h*(***x***) = *e*_*m*_ −*e* ≥ 0, we use the formulation of the Pontryagin maximum principle (PMP) described by Hartl, Sethi and Vickson [13]. Let *λ*_0_ ≥ 0 be the cost multiplier and ***λ*** = (*λ*_*v*_, *λ*_*e*_, *λ*_*l*_) the costate variable, and define the Hamiltonian

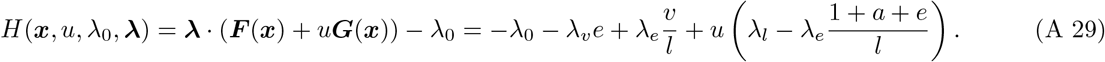

The derivative of the constraint function along the controlled dynamics is

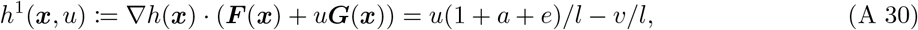

and we define the restricted control set

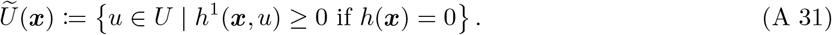

The PMP implies that, given an optimal control *u*^*∗*^(*t*) and corresponding trajectory ***x***^*∗*^(*t*), there exists an associated nowhere-vanishing costate trajectory ***λ***(*t*) such that, for almost every *t*, the function *u* ↦ *H*(***x***^*∗*^(*t*), *u, λ*_0_, ***λ***(*t*)) attains its maximum over 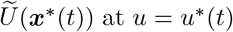. We distinguish between boundary arcs, where the constraint *h*(***x***) ≥ 0 is active, and interior arcs, where it is inactive. On an interior arc, the costate trajectory satisfies the adjoint equation

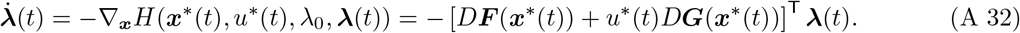

As noted in the main text, a key role is played by the switching function, equation (2.14). On an interior arc, the Hamiltonian maximization condition implies that the control *u*^*∗*^ is bang–bang unless *φ* vanishes on a nontrivial interval; such singular arcs are ruled out in the following section. On a boundary arc, *h* = 0 throughout the arc and hence *h*^1^ = 0, which requires *u*^*∗*^(*t*) = *v*^*∗*^(*t*)*/*(1 + *a* + *e*_*m*_).

Finally, we note that the terminal arc must have *u*^*∗*^ = *u*_*m*_. Since it ends at *e* = 0, it must be an interior arc and hence a bang arc, as singular arcs have been excluded. An arc with *u*^*∗*^ = 0 which starts outside the ray of fixed points E cannot reach it, since *V* is constant. Therefore, the terminal arc must have *u*^*∗*^ = *u*_*m*_.

#### (ii) Exclusion of singular arcs

A singular arc is an interior arc on which the switching function *φ* vanishes identically. We rule out such arcs using a standard argument involving Lie brackets of ***F*** and ***G***; see, for example, [17, Example 12.6]. Combining the state equation with the adjoint equation (A 32) gives the standard expressions^2^

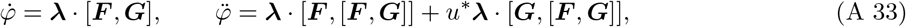

where the vector fields are evaluated at ***x***^*∗*^(*t*) and the costate at ***λ***(*t*). Direct calculation gives

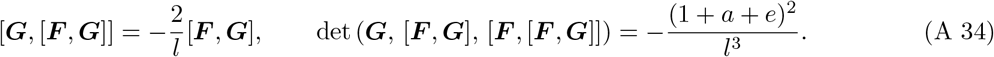

The determinant is nonzero throughout the taut region. Thus [***G***, [***F***, ***G***]] lies in the span of ***G*** and [***F***, ***G***], while ***G***, [***F***, ***G***], and [***F***, [***F***, ***G***]] are linearly independent. These conditions imply that the zeros of *φ* are isolated. In particular, *φ* cannot vanish on a nontrivial interval, so singular arcs do not occur and the optimal control is bang–bang on interior arcs.

#### (iii) Parameter-space boundaries

We derive the small-*v*_0_ asymptotic expressions for the region boundaries in figure 4*b*. For convenience, introduce the scaled parameters

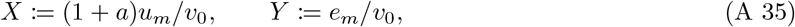

which are the axes of the diagram. We consider the limit *v*_0_ → 0 with *X, Y*, and *a* fixed. For the relevant trajectories, *v, e*, and *u* are *O*(*v*_0_), which also implies that *l* = 1 + *O*(*v*_0_) over time intervals of length *O*(1). Thus, to leading order, the dynamics reduce to

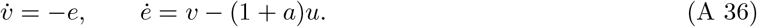

First consider the infeasible boundary. It is clear that the problem is feasible only if the control can be chosen, subject to 0 ≤*u* ≤*u*_*m*_, so that the first local maximum *e*_*p*_ of *e* satisfies *e*_*p*_ ≤*e*_*m*_. To see that the converse is true, note that after the first local maximum is reached, the trajectory can be continued using the proportional feedback control of section 2(b), with gain *γ* = (1 + *a*)*u*_*m*_*/*(2*e*_*p*_) chosen so that *u* ≤*u*_*m*_ for *e* ≤*e*_*p*_. Using an energy argument based on *V* (***x***) = (*le*^2^ + *v*^2^)*/*2, one can show that *e* does not subsequently exceed *e*_*p*_. By proposition 1, the feedback trajectory converges to *E*. Once sufficiently close, *E* can be reached in finite time by a *u* = 0 arc followed by a *u* = *u*_*m*_ arc.

It remains to determine the minimum possible *e*_p_. First assume 0 *< X <* 1 and define *Q* = (*v*− *Xv*_0_)^2^ + *e*^2^. For the leading-order dynamics (A 36) we have 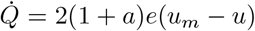, which is nonnegative while *e* ≥ 0. Let *t*^*′*^ be the first time at which *v* = *Xv*_0_. For *t < t*^*′*^,

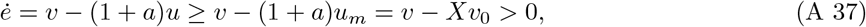

so the first local maximum of *e* cannot occur before *t*^*′*^. At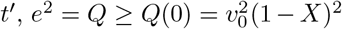, so *e*_*p*_ ≥*v*_0_(1 −*X*). This bound is sharp, since it can be attained by taking *u* = *u*_*m*_ for 0 ≤*t* ≤*t*^*′*^. For *X*≥ 1, the problem is feasible for any *Y >* 0: choose 0 *< e*_*p*_ *< e*_*m*_, take *u* = 0 until *e* = *e*_*p*_, and then apply the feedback control above. Since *v* ≤*v*_0_≤ (1 + *a*)*u*_*m*_, *e*_*p*_ is then a local maximum, after which the previous argument applies. Hence the infeasible boundary is

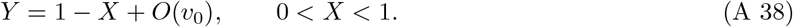

Next we consider the boundary between regions I and III, which occurs where *u*_*m*_ becomes too small for a one-switch bang–bang trajectory to reach *E*. At leading order, the initial *u* = 0 arc lies on the circle 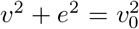. A terminal *u* = *u*_*m*_ arc that reaches (*v, e*) = (0, 0) lies on a circle centered at (*v*_0_*X*, 0) with radius *v*_0_*X*. A one-switch bang–bang trajectory exists if and only if these two circles intersect, which requires *X* ≥ 1*/*2. At *X* = 1*/*2, the circles are tangent at the initial point and the initial *u* = 0 arc vanishes. Hence the I/III boundary satisfies *X* = 1*/*2 + *O*(*v*_0_). The boundary is independent of *Y*, since the strain constraint is inactive in both regions I and III.

The remaining boundary of region I (the orange curve in figure 4*b*) indicates where the one-switch bang–bang trajectory violates the strain constraint. We find it by computing the maximum strain along this trajectory. Let (*v*_*s*_, *e*_*s*_) denote the switching point between the initial *u* = 0 arc and the terminal *u* = *u*_*m*_ arc. As noted above, the two arcs lie on circles centered at (0, 0) and (*v*_0_*X*, 0), respectively; computing their intersection yields *v*_*s*_*/v*_0_ = 1*/*(2*X*) and 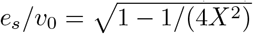. The terminal arc traverses its circle counterclockwise toward the target. If the switching point lies to the right of the top of this circle, (*v, e*) = (*Xv*_0_, *Xv*_0_), the strain continues to increase after the switch and reaches its maximum *Xv*_0_ at the top. If the switching point lies to the left of the top, the strain decreases immediately after the switch, so its maximum is *e*_*s*_. Since *v*_*s*_*/v*_0_ = 1*/*(2*X*), the transition occurs at 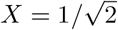. Thus, to leading order, the remaining boundary of region I is

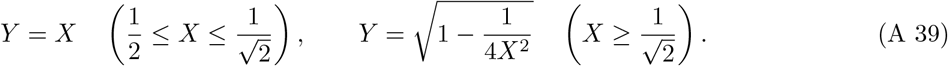

The first branch separates regions I and IV, while the second separates regions I and II.

Finally, consider the boundary between regions II and IV. On the boundary arc, *u* = *v/*(1 + *a* + *e*_*m*_), and since 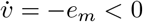, the required spinning rate is largest when the boundary is first reached. Hence the II/IV boundary is given exactly by equality in equation (2.13). In terms of the scaled parameters (A 35), this gives

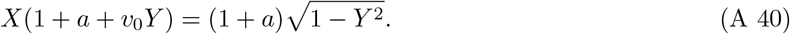

In the small-*v*_0_ limit, this reduces to *X*^2^ + *Y* ^2^ = 1 for 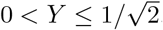.

### B. Analysis of the jumping task

#### (a) Estimate of aerodynamic rotational damping

To estimate the effect of air resistance on the spider’s rotation during a jump, we neglect the silk force and use a simple distributed drag model. We approximate the body as a uniform ellipsoid with longitudinal diameter *L* and transverse diameter *H*. For the drag calculation, we make the further approximation that its effective projected width is a constant *W* along the body. Let *ξ*∈ [−*L/*2, *L/*2] be a coordinate along the body axis, measured from the center of mass. We write *V >* 0 for a characteristic translational velocity normal to the body axis and 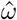 for the (dimensional) angular velocity. We assume 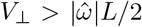, so that the local normal velocity 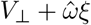 is positive along the entire body. Assuming quadratic pressure drag, the contribution from an element of length d*ξ* to the aerodynamic moment about the center of mass is

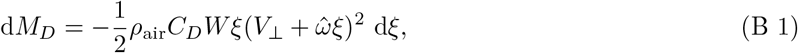

where *ρ*_air_ is the density of air and *C*_*D*_ is the drag coefficient. The terms proportional to 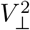 and 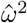 produce no net moment about the center of mass by symmetry. Integrating the remaining term over the body gives the aerodynamic damping moment *M*_*D*_:

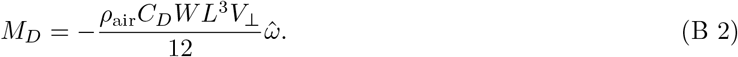

Treating *V*_*⊥*_ as constant over the flight and using 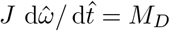, we obtain

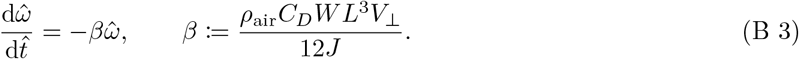

Let 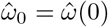. Over a flight of duration 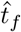, the angular displacement in the presence of aerodynamic damping satisfies

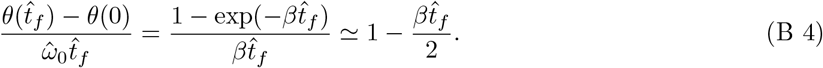

Here, we have normalized by 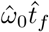, the angular displacement over the same duration in the absence of aerodynamic damping.

For the estimate, we take *d* = 3 mm, *m* = 1 · 10^*−*5^ kg, *L* = 2*d* = 6 mm, *W* = 5 mm, and 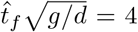.

Writing *J* = *jmd*^2^, as in section 3(a), the ellipsoidal approximation then gives

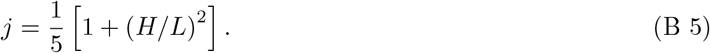

Thus *j* ≥ 0.2, with equality in the slender-body limit. Allowing for the legs and deviations from the ellipsoidal approximation, we take 0.2 ≤ *j* ≤ 0.5 as a plausible range. Based on the measured translational speeds in the jumping trials and the substantial body rotation during flight, we take *V*_*⊥*_ = 0.45 m*/*s. Taking 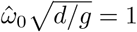 consistent with the observed values in figure 5, gives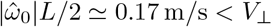.

Chen et al. [9] found that *C*_D_≃1.5 reproduced the observed deceleration of similarly sized salticids jumping without silk. Taking *ρ*_air_ = 1.2 kg*/*m^3^, the parameters above give 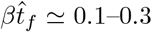. equation (B 4) then gives a reduction in angular displacement of roughly 5–15%. We therefore expect aerodynamic damping to have only a modest effect on the total body rotation during a jump.

### (b) Minimum strain and horizontal range near the zero-strain boundary

We analyze the terminal angle and horizontal range near the zero-strain boundary in figure 6, for target strains *ε*_*m*_ close to the minimum value 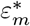 required for an upright landing. We work in nondimensional variables, using the notation of section 3(a). As in the figure, we take *x*_0_ = 1 and the launch angle to be *π/*4. For fixed *α* = *ω*_0_*/v*_0_, define 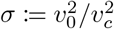, where *v*_*c*_ and *R*_*c*_ are defined in (3.8). The zero-strain boundary corresponds to *σ* = 1.

To determine the effect of a small target strain *ε*_*m*_ *>* 0, we consider a first-order perturbation about the ballistic (*ε*_*m*_ = 0) trajectory at the boundary *σ* = 1. Let 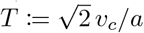 denote the flight time of the ballistic trajectory, and introduce the normalized time *τ* = *t/T*, so that *τ* = 0 at takeoff and *τ* = 1 at landing. At fixed *σ* = 1, we write

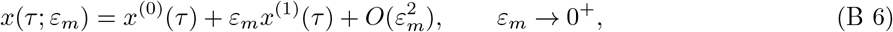

and similarly for *y, θ*, and the other state variables. We also expand the normalized flight time as 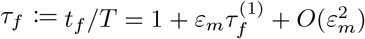. The zeroth-order term is precisely this ballistic reference trajectory:

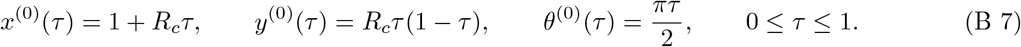

For *τ >* 0, let ***s***^(0)^(*τ*) denote the corresponding spinneret position and *ϕ*^(0)^(*τ*) = arg ***s***^(0)^(*τ*) the dragline angle. We also define the normalized switching time *τ*_*s*_ := *t*_*s*_*/T*, where *t*_*s*_ is given by (3.6). We assume 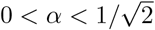 throughout, which is equivalent to the spinneret initially having positive vertical velocity. It then follows from (3.6) that *τ*_*s*_ *>*0. Moreover, *τ*_*s*_ *<* 1, since at landing ^(0)^(1) cos ^(0)^(1) − *x*^(0)^(1) sin *θ*^(0)^(1) = −(1 +*R*_c_) *<* 0. Since 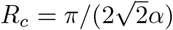, the same assumption on *α* implies *R*_*c*_ *> π/*2, a bound that will be used repeatedly below.

For *ε*_*m*_≪ 1, the loading interval *t*_*s*_ *< t < t*_*m*_ has duration *O*(*ε*_*m*_) and can be neglected to first order. We therefore approximate the strain as changing instantaneously from 0 to *ε*_*m*_ at *τ* = *τ*_*s*_. To express the resulting first-order perturbations at the unperturbed landing time, define

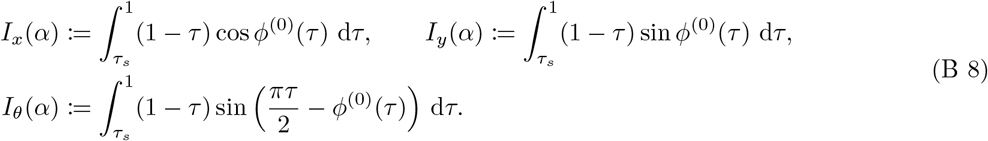

Taking the one-sided derivative of the equations of motion with respect to *ε*_*m*_ at *ε*_*m*_ = 0 and integrating twice from *τ*_*s*_ to 1 gives

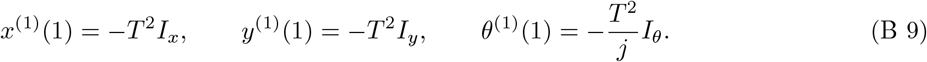

To obtain the first-order correction to the terminal angle and range, we must also account for the change in the landing time. Expanding the landing condition *y*(*τ*_*f*_; *ε*_*m*_) = 0 about *τ* = 1, using (B 7) and (B 9), gives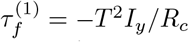. Substituting this correction into *θ*_*f*_ = *θ*(*τ*_*f*_) and *R* = *x*(*τ*_*f*_) − 1 then gives the first-order corrections

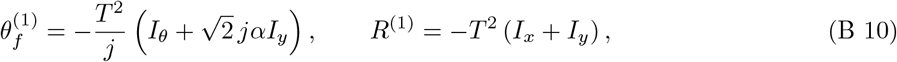

where we used the expression for *R*_*c*_ given above.

We now use these first-order corrections to determine how 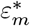 and the corresponding range vary as *σ* increases through the zero-strain boundary. For *ε*_*m*_ = 0, ballistic motion gives *θ*_*f*_ (*σ*, 0) = *πσ/*2 and *R*(*σ*, 0) = *R*_*c*_*σ*. For *σ >* 1, the minimum target strain satisfies 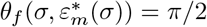 by definition. Differentiating this condition at *σ* = 1^+^ gives

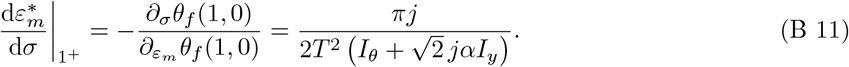

The corresponding horizontal range is 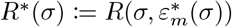. We normalize this by the range at the zero-strain boundary and define *ρ*(*σ*) := *R*^*∗*^(*σ*)*/R*_*c*_. Using the chain rule,

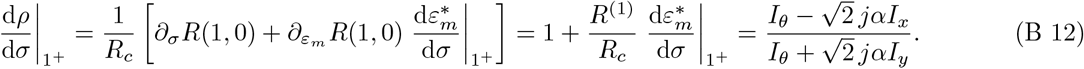

It remains to show that this derivative is positive for 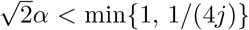. We sketch the proof, which proceeds in three steps. First, we derive bounds on *τ*_*s*_ and *ϕ*^(0)^. We then use these bounds to estimate the ratios *I*_*y*_*/I*_*x*_ and *I*_*θ*_*/I*_*x*_. Finally, we apply these estimates to the right-hand side of (B 12).

We first derive bounds on the switching time *τ*_*s*_ along the ballistic trajectory. Write ***s***^(0)^(*τ*) = (*ξ*(*τ*), *η*(*τ*)) for the coordinates of the spinneret:

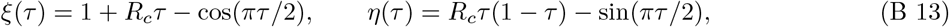

so that tan *ϕ*^(0)^(*τ*) = *η*(*τ*)*/ξ*(*τ*). The switching condition (3.6) along the ballistic trajectory is equivalent to *η*(*τ*_*s*_)*/ξ*(*τ*_*s*_) = tan (*πτ*_*s*_*/*2). Substituting the above expressions yields

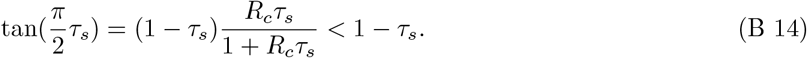

From this it follows that *τ*_*s*_ *<* 3*/*8: otherwise tan(*πτ*_*s*_*/*2) ≥tan(3*π/*16) *>* 5*/*8, while 1− *τ*_*s*_ ≤5*/*8, contradicting (B 14).

Next, we need the following bounds on *ϕ*^(0)^ for *τ*_*s*_ ≤ *τ* ≤ 1:

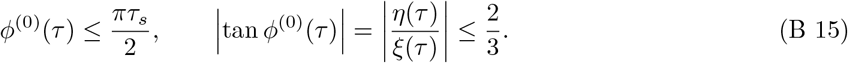

Define *p*(*τ*) := *η*(*τ*) cos(*πτ*_*s*_*/*2) − *ξ*(*τ*) sin(*πτ*_*s*_*/*2); then the first bound is equivalent to *p*(*τ*) ≤ 0. It follows immediately from the switching condition that *p*(*τ*_*s*_) = 0. Direct differentiation and the inequality (B 14) give *p*^*′*^(*τ*_*s*_) *<* 0, while 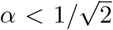 together with *τ*_*s*_ *<* 3*/*8 gives *p*^*′′*^(*τ*) *<* 0 for *τ*_*s*_≤ *τ*≤ 1. Hence *p*(*τ*) ≤0 throughout this interval, which proves the first bound. The upper bound *η/ ξ* < 2*/*3 follows easily: if *τ*_*s*_ ≤ 1*/*3, then 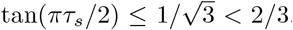, while if *τ*_*s*_ *>* 1*/*3, (B 14) gives tan(*πτ*_*s*_*/*2) *<* 1 − *τ*_*s*_ *<* 2*/*3. In either case, *η/ξ* ≤ tan(*πτ*_*s*_*/*2) *<* 2*/*3. For the lower bound, define *q*(*τ*) := *η*(*τ*) +(2*/*3)*ξ*(*τ*). Then *q*(0) = 0, while *R*_*c*_ *> π/*2 gives *q*(1) = (2*R*_*c*_ − 1)*/*3 *>* 0. Moreover,

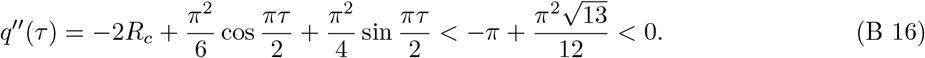

Thus *q* is concave and lies above the chord joining its endpoint values, so *q*(*τ*) ≥ 0 for 0 ≤ *τ* ≤ 1. Hence *η/ξ* ≥ −2*/*3, completing the proof of (B 15).

We now use (B 15) to estimate the ratios *I*_*y*_*/I*_*x*_ and *I*_*θ*_*/I*_*x*_. The second bound gives 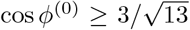. Rewriting sin *ϕ*^(0)^ = tan *ϕ*^(0)^ cos *ϕ*^(0)^ in the definition of *I*_*y*_, together with the same bound, gives |*I*_*y*_| ≤ (2*/*3)*I*_*x*_. For *I*_*θ*_*/I*_*x*_, the first bound in (B 15) gives

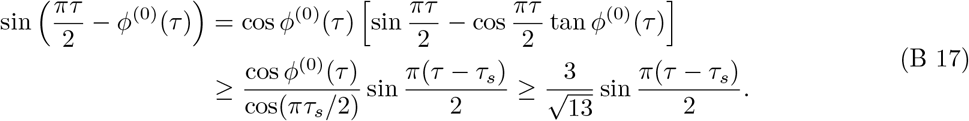

Since *I*_*x*_ ≤ (1 − *τ*_*s*_)^2^*/*2, setting *z* := (*π/*2)(1 − *τ*_*s*_) gives

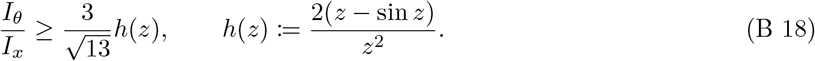

Direct differentiation shows that *h* is increasing for 0 *< z < π/*2, and since *τ*_*s*_ *<* 3*/*8,

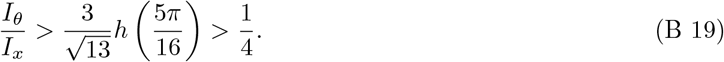

Together with |*I*_*y*_| ≤ (2*/* 3) *I*_*x*_, this shows that both the numerator and denominator in (B 12) are positive provided 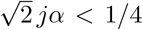. Hence the derivative in (B 12) is positive whenever 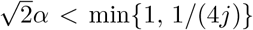. For *j* ≤ 1*/*4, the derivative is positive throughout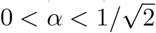.

## Footnotes

1 For *x*_0_ = 1, the strain is formally undefined at takeoff; we understand this case as the limit *x*_0_ = 1 + *δ* as *δ* → 0^+^.

2 We use the convention [***X, Y***] = *D****Y X*** *− D****X Y***.

